# LOXL2-dependent deacetylation of aldolase A induces metabolic reprogramming and tumor progression

**DOI:** 10.1101/2022.03.06.483165

**Authors:** Ji-Wei Jiao, Xiu-Hui Zhan, Juan-Juan Wang, Li-Xia He, Zhen-Chang Guo, Xiu-E Xu, Lian-Di Liao, Xin Huang, Bing Wen, Yi-Wei Xu, Hai Hu, Gera Neufeld, Zhi-Jie Chang, Kai Zhang, Li-Yan Xu, En-Min Li

## Abstract

Lysyl-oxidase like-2 (LOXL2) regulates extracellular matrix remodeling and promotes tumor invasion and metastasis. Altered metabolism is a core hallmark of cancer, however, it remains unclear whether and how LOXL2 contributes to tumor metabolism. Here, we found that LOXL2 and its catalytically inactive L2Δ13 splice variant also function as novel deacetylases that trigger metabolic reprogramming during malignant transformation. Integrated transcriptomic and metabolomic analysis revealed that L2Δ13-overexpressing transgenic mice displayed perturbed glucose and lipid metabolism, which was associated with increased hepatic fibrosis and enhanced formation of precancerous lesions induced by chemical carcinogens, such as carbon tetrachloride and N-nitrosomethylbenzylamine. Furthermore, both LOXL2 and L2Δ13 boosted glucose metabolism of esophageal tumor cells, thereby facilitating tumor cell proliferation *in vitro* and *in vivo.* Mechanistically, LOXL2 and L2Δ13 interacted physically with several glycolic proteins including aldolase A to enhance their enzymatic activities and mobilization from the actin cytoskeleton. Using SILAC followed by proteomic analysis, we identified LOXL2 as a deacetylase targeting metabolic proteins in esophageal cancer. Importantly, both LOXL2 and L2Δ13 directly catalyzed the deacetylation of aldolase A at K13, resulting in enhanced glycolysis which subsequently reprogramed tumor metabolism and promoted tumor progression. High level expression of LOXL2/L2Δ13 combined with decreased acetylation of aldolase-K13 predicted poor clinical outcome in patients with esophageal cancer. In summary, we have characterized a novel molecular mechanism that mediates the pro-tumorigenic activity of LOXL2 independently of its classical amine oxidase activity. These findings may enable the future development of therapeutic agents targeting the metabolic machinery via LOXL2 or L2Δ13.

## Introduction

The lysyl oxidase (LOX) family is comprised of five members (LOX and LOXL1-4) whose primary function is to oxidize the ε-amino group of peptidyl lysine residues to aldehyde residues, thereby covalently crosslinking elastin and/or collagens in the extracellular matrix (ECM) (1,2). The ECM remodeling provides structural scaffolding and contextual information for cells, which is required for both tissue integrity and expansion. Aberrant expression and activity of lysyl oxidase-like 2 (LOXL2) as a secreted copper- and quinone-dependent enzyme is involved in pathological processes predominantly associated with the ECM remodeling, such as Wilson’s disease, heart failure and cholestasis (3–5). LOXL2-induced alterations of ECM organization are associated with the induction of abnormal fibrosis, which in tumors promotes tumor cell invasiveness and tumor metastasis (6–8). Moreover, elevated expression of LOXL2 as a result of hypoxic stress has been found to modulate gene transcription and to promote epithelial-to-mesenchymal transition (EMT), contributing to the enhancement of tumor cell motility, invasiveness and metastasis (9,10). It has also been reported that specific catalytically inactive mutants of LOXL2 mediate keratinocyte differentiation and EMT caused by an interaction with the transcription factor Snail1 (11–13). However, the contribution of the non-enzymatic functions of LOXL2 to tumor progression has not yet been studied in depth.

Changes in the organization of the ECM can influence cellular metabolism (14). For example, degradation of ECM-associated hyaluronic acid can affect glucose uptake and enhance glycolysis (15). LOXL2 is also present in intra-cellular compartments, such as the nucleus and cytoplasm of cells (9,16,17), suggesting that LOXL2 as well as additional lysyl oxidases perhaps also affects metabolism through interacting with intracellular proteins. Although elevated expression of lysyl oxidases and metabolic reprogramming are coincident in multiple biological contexts (18,19), their underlying mechanistic links are not well established.

We have previously identified a novel LOXL2 splice variant L2Δ13 in human esophageal squamous cell carcinoma cells (mRNA, GenBank accession number KF928961; protein, GenBank accession number AHJ59530.1) (20). The L2Δ13 isoform was also observed in immortalized esophageal epithelial cells and in various types of cancer cells (20). Unlike extracellular full-length LOXL2, L2Δ13 lacks exon 13, which is located in the highly conserved catalytic C-terminal domain of LOXL2, resulting in the complete loss of amine oxidase activity. In addition, L2Δ13 fails to be secreted from cells. Thus, L2Δ13 represents a LOXL2 form that promotes certain biological activities independent of the classical enzymatic function of the lysyl oxidases. However, some of these enzyme-independent functions of L2Δ13 are likely to be shared with full-length LOXL2. High level expression of full-length LOXL2, as well as its L2Δ13 isoform, promotes cell migration and invasion of esophageal cancer cells *in vitro* and *in vivo,* which is further linked to tumor metastasis and poor clinical outcome of esophageal cancer patients (16,21). In this study, we have generated for the first time a genetic mouse model overexpressing L2Δ13 to systematically explore the *in vivo* biological contributions of L2Δ13 ranging from its physiological functions to its role in malignant transformation. In particular, L2Δ13 and LOXL2 physically interact with and simultaneously deacetylate glycolic enzymes, such as aldolase, to accelerate glucose metabolism, suggesting that LOXL2/L2Δ13 induces metabolic alterations which in turn may be responsible for part of the pro-tumorigenic effects of LOXL2.

## Materials and Methods

### Mouse models

For induction of hepatic fibrosis by carbon tetrachloride (CCl_4_), 8-week-old L2Δ13-overexpressing mice (Δ13/Δ13) and wild-type mice received intraperitoneal injections of CCl_4_ (2 mL/kg; Aladdin) dissolved in corn oil at a ratio of 1:4 or corn oil alone (2 mL/kg; Aladdin) twice a week for 6 weeks. At the end of the experiments, mice were sacrificed and livers were removed for further analysis. Necrosis was assessed by morphometric analysis. Blood samples were quickly collected, and sera were harvested by centrifugation for blood biochemical analysis.

For induction of esophageal lesions by the nitrosamine compound N-nitrosomethylbenzylamine (NMBA), 8-week-old Δ13/Δ13 and wild-type mice were injected subcutaneously with NMBA (Wuhan 3B Scientific Corporation) diluted in 20% dimethyl sulfoxide (DMSO), at a dose of 1 mg/kg, three times a week for 12 weeks. NMBA-treated mice were euthanized at 46 weeks of age and their esophagi were dissected and analyzed. Animal studies were conducted in accordance with protocols approved by the Animal Research Committee of the Shantou Administration Center.

### Patients and tissue specimens

Human esophageal carcinoma tissue specimens were collected directly after surgical resection between November 2007 and January 2011 at Shantou Central Hospital (Sun Yat-sen University, Shantou, China). The study was approved by institutional review board, and written informed consent was obtained from all patients.

### Metabolomics

Liver tissues were harvested from wild-type and L2Δ13-overexpressing mice, rinsed with saline solution and immediately transferred to a tube placed in liquid nitrogen. The frozen tissues were minced, weighed, and 25 mg was dissolved in an equal volume of water:methanol (1:1, v:v) solution. Samples were disrupted with a TissueLyser through high-speed shaking, precipitated overnight and centrifuged. Chromatographic separations of resulting supernatant of samples were performed using an ultra-performance liquid chromatography (UPLC) system (Waters). The injection volume for each sample was 10 μL. A high-resolution tandem mass spectrometer Xevo G2 XS QTOF (Waters) was subsequently applied to detect metabolites of eluted hepatic samples. Mass spectrometry data were acquired in the Centroid MSE mode and analyzed using MetaX. Differential metabolites with a fold change ≥ 1.2 and *q*-value < 0.05 were considered as significant. The metabolite profile was analyzed by the Pathway Analysis module of the MetaboAnalyst.

### Cell permeabilization, fractionation and aldolase activity assays

For aldolase activity in the diffusible fraction (in the supernatant), cells were seeded into 6-well plates and permeabilized with digitonin (30 μg/mL; Sigma-Aldrich), followed by the collection of supernatant and cell lysate for further analyses as described previously (22). For cell fractionation, ProteoExtract®Cytoskeleton Enrichment and Isolation Kit (Millipore) was applied to isolation of cytoskeletal and soluble compartments of indicated cells.

### *In vitro* deacetylation assay

Purified GST-tagged aldolase A (0.5 μg) or synthetic acetyl peptide of aldolase A-K13 (98% purity) was incubated with purified Flag-tagged LOXL2/L2Δ13 (0.25 μg) in a 50 nM ATP-containing reaction buffer at 30□ for 2 h. The buffer contains 50 mM Tris-HCl (pH 9.0), 50 mM NaCl, 4 mM MgCl_2_, 0.5 mM DTT, 0.2 mM phenylmethylsulfonyl fluoride, 0.02% Nonidet P-40 and 5% glycerol(23). The reaction mixtures were subjected to Western blotting or Dot blotting analysis.

### Statistical analyses

Statistical analyses were performed using GraphPad Prism 7 software (GraphPad) or SPSS 22.0 software (IBM). Unless otherwise stated, all experiments were repeated independently three times.

Comparative data were evaluated by an unpaired Student’s *t*-test or nonparametric test. Survival curves were estimated using the Kaplan-Meier method with log-rank test. Differences were considered statistically significant when the two-tailed *p*-value was < 0.05.

## Results

### The L2Δ13 spliced isoform of LOXL2 regulates metabolic reprogramming

Using a Cre/LoxP-based gene-targeting strategy, we generated a gain-of-function mouse model to explore the *in vivo* biological role of the human *L2Δ13* gene. Flag-tagged human *L2Δ13* cDNA was inserted by homologous recombination into the *ROSA26* locus and expressed under the control of the CAG promoter following Cre-mediated excision (Fig. 1A; Supplementary Fig. S1A). Targeted Flag-tagged L2Δ13 was localized in the cytoplasm of cells of different organs of homozygous L2Δ13-mice (Supplementary Fig. S1B and S1C). It has been reported that overexpression of fulllength LOXL2 results in severe male sterility (10). However, L2Δ13-overexpressing male homozygous mice from the same genetic background did not display male sterility, which was further verified by hematoxylin and eosin (H&E) staining of their testes and epididymides (Supplementary Fig. S1D). Interestingly, body weights of homozygous L2Δ13-overexpressing mice were significantly reduced compared with both littermate wild-type and heterozygous L2Δ13-overexpressing mice with almost equivalent food intake and water intake (Fig. 1B; Supplementary Fig. S1E and S1F). The lower body mass of the L2Δ13 mice was accompanied by decreased body fat and lower blood glucose with age (Fig. 1C and 1D; Supplementary Table S1). In contrast, serum cholesterol (CHOL), low density lipoprotein cholesterol (LDL) and triglyceride (TG) levels were greatly increased in the L2Δ13 mice compared with age-matched wild-type mice (Fig. 1D; Supplementary Table S1). Oil Red O staining further indicated that the percentage of mice with massive lipid deposition in the liver of L2Δ13-overexpressing mice was much higher than that of wild-type mice (Fig. 1E). Moreover, overexpression of L2Δ13 did not interfere with kidney function indexes, including blood urea nitrogen, creatinine and uric acid (Supplementary Table S1). These findings imply that the reduced body weight is due in part to liver injury that may be caused by modified lipid metabolism. To further examine the involvement of L2Δ13 in metabolic regulation, we fed L2Δ13-overexpressing mice and wild-type control mice with either a normal chow diet (NCD) or high-fat diet (HFD) for eight weeks (Fig. 1F). The wild-type mice gradually gained body weight and displayed a higher percentage of body fat following HFD, whereas L2Δ13-overexpressing mice failed to gain body weight at a similar pace (Fig. 1G; Supplementary Fig. S1G). Notably, L2Δ13 overexpression resulted in reduced hepatic glycogen stores in mice that were maintained on the HFD diet, suggesting that L2Δ13 enhances glucose consumption (Fig. 1H). The L2Δ13-overexpressing mice had lower concentrations of glucose in their sera, while the concentration of cholesterol in their sera was higher than that observed in wild-type mice on HFD diet (Fig. 1I). Taken together, these observations suggest that L2Δ13 functions as a regulator of metabolic carbon balance.

**Figure 1.**
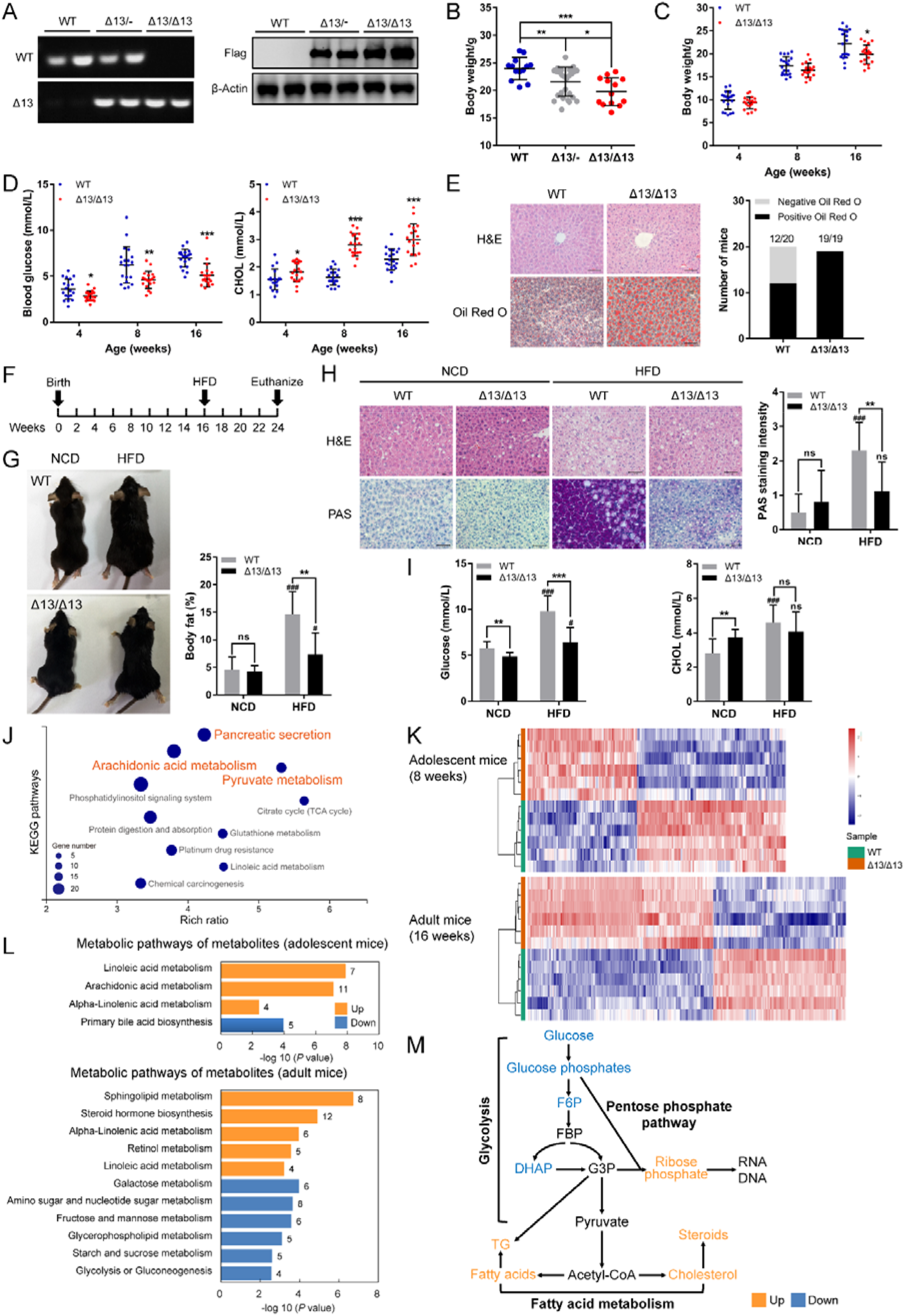
L2Δ13 overexpression leads to metabolic imbalance. **(A)** Left: PCR conformation of overexpression of L2Δ13 in C57BL/6J mice; right: western blotting analysis of livers from wild-type (WT), heterozygous (Δ13/-) and homozygous L2Δ13-mice (Δ13/Δ13). **(B)** Body weight of male littermate mice at 8 weeks of age (n = 13, 28, 13). **(C and D)** Body mass **(C)**, blood glucose concentrations and CHOL concentrations **(D)** from wild-type and L2Δ13-overexpressing mice at 4, 8 and 16 weeks of age (n = 19 or 20). **(E)** Left: representative H&E and Oil Red O staining of livers from wild-type and L2Δ13-overexpressing mice; right: number of mice with positive Oil Red O staining. **(F)** Schematic diagram of Δ13/Δ13 mice and littermate wild-type controls fed either a normal chow diet (NCD) or high-fat diet (HFD) for 8 weeks (n = 8-10). **(G)** Images (left) and body fat percentage (right) of mice from **(F). (H)** H&E and PAS staining of livers from mice. **(I)** Blood glucose concentrations and CHOL concentrations of the mice from **(G)**. **(J)** Significant patterns for KEGG pathways of differentially-expressed genes of L2Δ13-overexpressing and age-matched wild-type mice (n = 6; GSE145238). (K) Heatmap of metabolite clusters in L2Δ13-overexpressing and paired wild-type mice measured by LC-MS-based metabolomics (n = 6). **(L)** Metabolic pathways analysis of the significantly enriched metabolites (*q* value < 0.05, fold change ≥ 1.2) using MetaboAnalyst. **(M)** Schematic representation of the L2Δ13-regulated metabolic flux in mice under physiological conditions. F6P, fructose-6-phosphate; FBP, fructose-1,6-bisphosphate; G3P, glyceraldehyde 3-phosphate; Actyl-CoA, actyl coenzyme A. **(B-E, G-I)** Bar graphs represent means ± SD. *P*-value was calculated by *t*-test. * *P* < 0.05, ** *P* < 0.01 or *** *P* < 0.001. ^#^ Different from groups of mice fed NCD, ^#^*P* < 0.05, ^###^*P* < 0.001. ns: not significant. Scale bar, 50 μm.

To determine which genes are regulated by L2Δ13 *in vivo*, next generation RNA sequencing was performed to compare the RNA expression profile of liver tissues excised from L2Δ13-overexpressing and wild-type mice (GSE145238). Transcriptomic data identified 599 genes differentially expressed by greater than a 2-fold change in L2Δ13-overexpressing mice relative to paired wild-type samples (Supplementary Fig. S1H). Importantly, more than 60% of these highly altered genes were predominantly involved in diverse metabolic processes, including pyruvate metabolism, TCA cycle, lipid metabolism and amino acid metabolism (Fig. 1J). To determine if these L2Δ13-induced changes in gene expression are reflected by changes in the concentration of metabolites, we measured the levels of metabolites contained in mouse livers from these two groups by liquid chromatography-mass spectrometry (LC-MS) analysis. Consistently, metabolomics data derived from the livers of both adolescent and adult mice showed significant differences in the levels of metabolites between L2Δ13 mice and wild-type mice, further indicating that L2Δ13 induces metabolic reprogramming (Fig. 1K). Metabolomic pathway analysis revealed that L2Δ13 accelerates glycolysis/gluconeogenesis, fructose and mannose metabolism, and amino acid metabolism (Fig. 1L; Supplementary Fig. S1I). Moreover, L2Δ13 contributed to higher downstream lipid metabolism, as illustrated by enrichments in intermediate metabolites of linoleic acid metabolism, sphingolipid metabolism, as well as steroid hormone biosynthesis (Fig. 1L; Supplementary Fig. S1I). These results indicate that L2Δ13 overexpression enhances the expression of enzymes important for glucose consumption and lipid metabolism (Fig. 1M), suggesting that L2Δ13 may also be able to induce metabolic reprogramming under physiological conditions.

### Full-length LOXL2 and L2Δ13 enhances tumor cell proliferation and the formation of precancerous lesions

Carbon tetrachloride (CCl_4_) is quickly metabolized to initiate a lipid peroxidation chain reaction in the liver, thereby inducing both acute and chronic liver injury (24,25). To assess whether L2Δ13 affects the development of liver fibrosis, L2Δ13-overexpressing and wild-type mice were intraperitoneally injected with hepatotoxic CCl_4_ twice a week for six weeks (Fig. 2A). Importantly, overexpression of L2Δ13 promoted fibrotic nodule formation and focal necrosis, relative to similarly treated wild-type mice following CCl_4_ treatment (Fig. 2B and 2C). In mice overexpressing L2Δ13, we also found increased levels of serum alanine aminotransferase (ALT) following exposure to CCl_4_ as compared with wild-type mice exposed to CCl_4_ (Fig. 2D). Furthermore, L2Δ13-overexpressing mice contained a dramatically higher concentration of α-smooth muscle actin (α-SMA)-positive cells and sirius red-positive staining of collagen in their livers as compared with control wild-type mice, indicating that L2Δ13 overexpression aggravates CCl4-induced liver injury by enhancement of liver fibrosis, even though L2Δ13 is not secreted and lacks enzyme activity, and thus cannot affect fibrosis directly (Fig. 2E and 2F).

**Figure 2.**
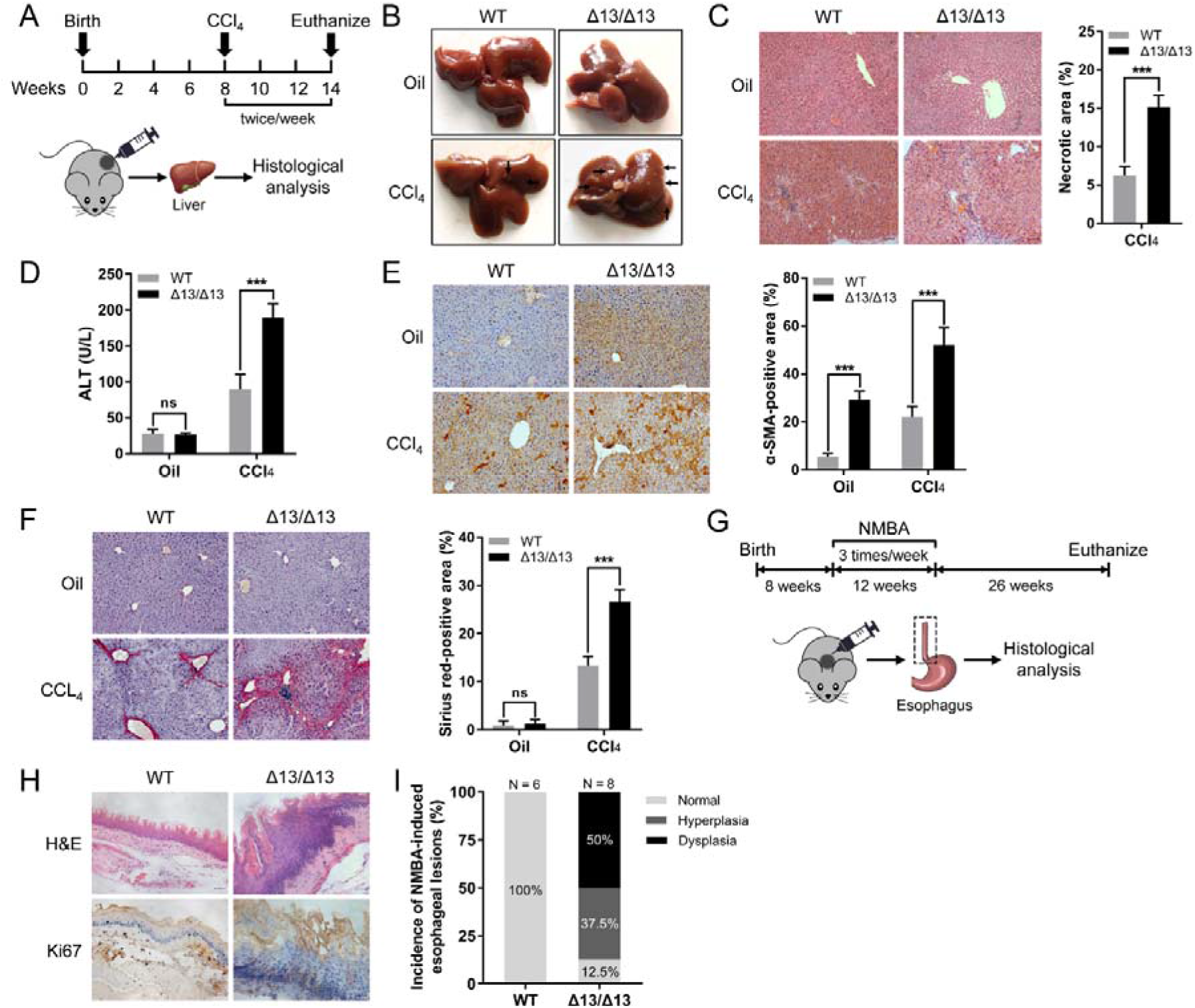
L2Δ13 enhances hepatic fibrogenesis and esophageal lesions. **(A)** Schematic diagram of CCl_4_-induced liver fibrosis in mice. Eight-week-old wild-type and L2Δ13-overexpressing mice were intraperitoneally injected with either CCl_4_ (2 mL/kg) dissolved in corn oil or corn oil alone (2 mL/kg) twice a week for 6 weeks. CCl_4_, carbon tetrachloride. **(B)** Representative images of livers from WT and Δ13/Δ13 mice (top: oil-injected, n = 6-8; bottom: CCl4-injected, n = 12-16). Arrows indicate fibrotic nodules visible on CCl4-treated mice. **(C)** H&E and percentage of focal necrosis area of livers from mice. Scale bar, 50 μm. **(D)** Sera from mice in (B) were harvested for serum alanine aminotransferase (ALT) analysis. **(E and F)** Left: representative images of α-SMA staining or sirius red staining of liver sections from **(C)**; right, quantification of α-SMA-positive or sirius red-positive areas. Scale bar, 50 μm. Bar graphs of **(C-F)** represent means ± SD. *** *P* < 0.001 by *t*-test analysis. ns: not significant. **(G)** Schematic overview of NMBA-induced esophageal lesions in mice. Eight-week-old wild-type and L2Δ13-overexpressing mice received subcutaneous injections of NMBA in 20% DMSO (1 mg/kg) three times a week for 12 weeks, and were euthanized at the age of 46 weeks (n = 6, 8). **(H)** Representative paraffin sections stained with H&E and anti-Ki67 antibody (a marker of cell proliferation) for esophageal tissues derived from wild-type and Δ13/Δ13 mice treated with NMBA. **(I)** Percentage of incidence of NMBA-induced esophageal lesions.

Changes in metabolism are known to be associated with the tumorigenic transformation of cells (26), and we therefore wondered if L2Δ13 could promote malignant transformation induced by carcinogens. We administered subcutaneously nitrosamine compound N-nitrosomethylbenzylamine (NMBA), a known carcinogen that specifically induces esophageal cancer, to L2Δ13-overexpressing and wild-type mice three times a week for 12 weeks (Fig. 2G) (27). Histological analysis revealed that L2Δ13 mice more readily developed NMBA-induced precancerous lesions in the esophageal squamous epithelium, as evidenced by cellular proliferation and dysplasia, relative to wild-type mice treated similarly (Fig. 2H). Compared with wild-type mice exposed to NMBA, the incidence of development and degree of esophageal lesions were strongly enhanced in L2Δ13-overexpressing mice (Fig. 2I), suggesting that L2Δ13 has an additive or synergistic effect with NMBA that leads to enhanced formation of precancerous esophageal lesions.

A high proportion of patients with L2Δ13-positive tumor cells existed in various types of malignant cancer (Fig. S2). Previous studies have linked altered metabolism with cell growth and proliferation in cancer cells (28). However, the contribution of either full-length LOXL2 or its spliced isoforms to tumor cell proliferation remains unclear. We silenced LOXL2 expression in esophageal cancer cells using specific siRNAs or shRNAs, and then conducted rescue experiments by ectopically re-expressing full-length LOXL2 or L2Δ13 in the LOXL2-silenced cancer cells as previously described (Fig. 3A; Supplementary Fig. S3A) (16). LOXL2 depletion strikingly inhibited both the proliferation and metabolism of the cells, as determined by the measurement of DNA synthesis and by the accumulation of neutral lipids (Fig. 3B and 3C; Supplementary Fig. S3B-S3D). In contrast, re-expression of either LOXL2 or L2Δ13 in LOXL2-depleted cells fully rescued their impaired proliferative capacity (Fig. 3D and 3E). Surprisingly, L2Δ13 seemed more effective than full-length LOXL2 in the colony formation assay (Fig. 3E). These *in vitro* results were further verified by *in vivo* xenograft rescue experiments. The development of subcutaneous tumors from implanted KYSE510 cells silenced for LOXL2 expression was strongly inhibited, whereas re-expression of LOXL2 or L2Δ13 in the LOXL2-silenced cells partially restored tumor development (Fig. 3F and 3G; Supplementary Fig. S3E). Interestingly, L2Δ13 in this assay also rescued tumor development more efficiently than full-length LOXL2.

**Figure 3.**
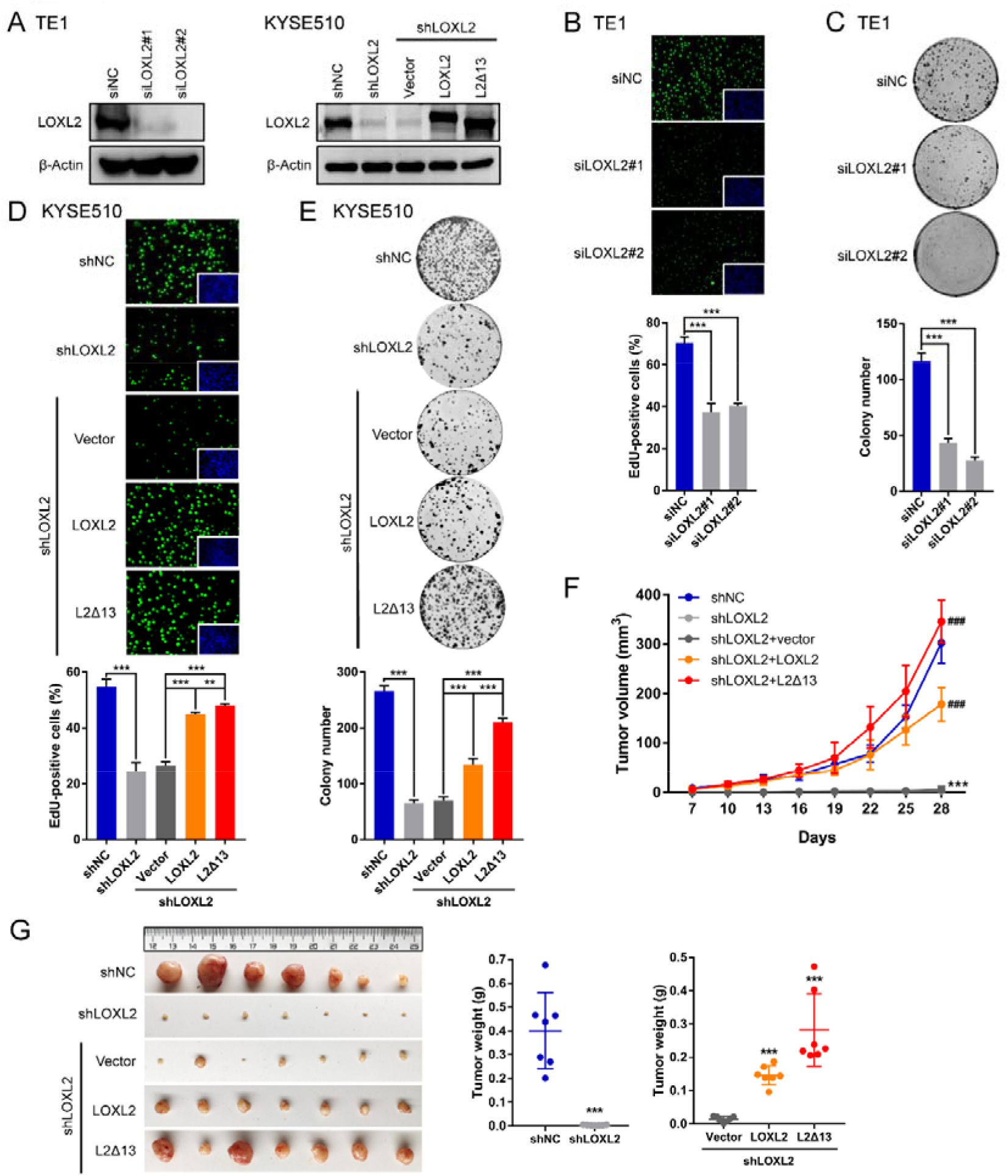
Full-length LOXL2 and L2Δ13 promote tumor cell proliferation *in vitro* and *in vivo*. **(A)** Western blotting assays of esophageal cancer cells following depletion of LOXL2 by specific siRNAs (left) and LOXL2 stably-depleted esophageal cancer cells following ectopic re-expression of LOXL2 or L2Δ13 (right). **(B and C)** EdU **(B)** and colony formation **(C)** assays of esophageal cancer cells following silencing of LOXL2. Error bars indicate mean ± SD; n = 5 for EdU, n = 3 for colony formation. \*\*\**P* < 0.001. **(D and E)** EdU **(D)** and colony formation **(E)** assays for KYSE510 cells infected with a scrambled shRNA or shLOXL2 and LOXL2-silenced KYSE510 cells expressing empty vector, LOXL2-Flag or L2Δ13-Flag. Error bars represent mean ± SD of triplicates. \*\**P* < 0.01 or \*\*\**P* < 0.001. **(F)** Indicated KYSE510 stably-infected cells (1 × 10^6^ in 100 μL PBS) were implanted subcutaneously into nude mice. Tumor volumes of xenograft mice were measured at the indicated time points, and tumors were excised after 30 days (n = 7). *Different from xenografts injected KYSE510 cells infected with shNC, \*\*\**P* < 0.001; ^#^different from xenografts injected with LOXL2-depleted cells expressing vector, ^###^*P* < 0.001. *P*-values were determined by a *t*-test. **(G)** Images of dissected tumors and summary of average tumor weights measured at end point.

### LOXL2 and L2Δ13 drive glycolysis by interacting with glycolic proteins in both nonmalignant and cancer cells

To characterize the molecular mechanisms by which LOXL2 regulates metabolism, we analyzed the metabolic status of esophageal nonmalignant cells and esophageal cancer cells in the presence and absence of LOXL2. Glycolysis, an essential pathway for glucose metabolism, utilizes glucose to generate pyruvate and subsequently ATP and lactate (29). Intriguingly, ATP levels, along with glucose uptake and lactate production, were enhanced in esophageal epithelial cells overexpressing either LOXL2 or L2Δ13, indicating that LOXL2 and L2Δ13 may accelerate metabolism via enhancement of glycolysis (Fig. 4A; Supplementary Fig. S4A). Glucose metabolism shifts from oxidative phosphorylation to aerobic glycolysis in proliferating cancer cells (the Warburg effect) (30). Consistently, depletion of LOXL2 greatly decreased ATP formation and lactate buildup as a result of reduced availability of glucose in TE1 and KYSE510 esophageal cancer cells (Fig. 4B; Supplementary Fig. S4B). ATP formation was completely rescued and lactate production increased to background levels following ectopic expression of LOXL2 or L2Δ13 (Fig. 4C), suggesting that LOXL2 and L2Δ13 specifically promote glycolysis in cancer cells. Furthermore, microarray data of patients with esophageal cancer also indicated that LOXL2 expression positively correlated with enhanced glycolysis and gluconeogenesis pathways in clinical settings (Fig. 4D).

**Figure 4.**
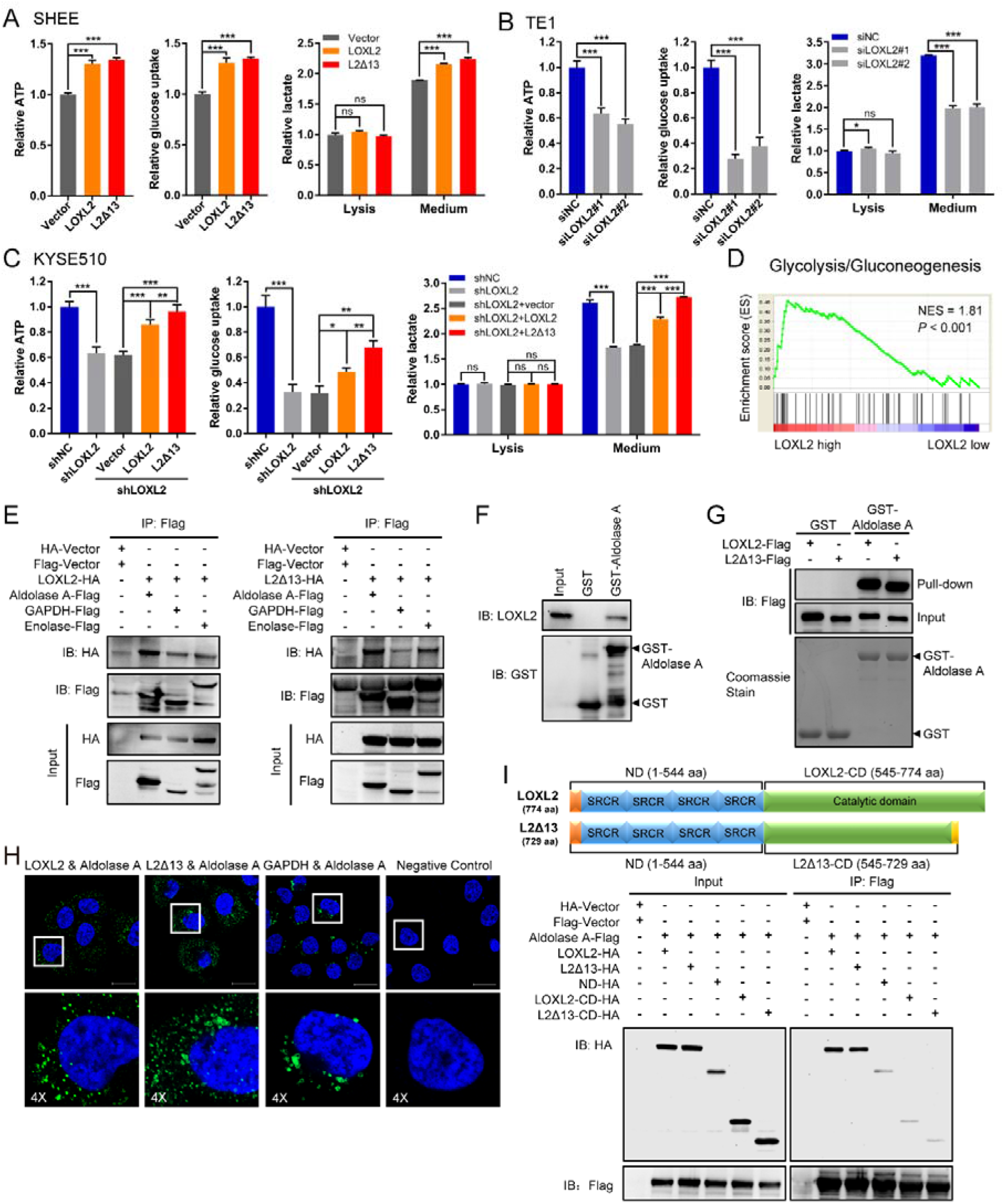
LOXL2 and L2Δ13 regulate glycolysis through interacting with glycolic enzymes. **(A-C)** Levels of ATP, glucose uptake and lactate of nonmalignant esophageal epithelial cells expressing recombinant full-length LOXL2, L2Δ13 variant or control vector **(A)**, esophageal cancer cells following depletion of LOXL2 by specific siRNAs **(B)** and LOXL2-depleted esophageal cancer cells following ectopic re-expression of LOXL2 or L2Δ13 **(C)**. Mean ± SD, n = 3-5. \**P* < 0.05, \*\**P* < 0.01 or \*\*\**P* < 0.001. ns, not significant. **(D)** Gene set enrichment analysis (GSEA) plot shows that LOXL2 gene expression positively correlated with genes in the glycolysis/gluconeogenesis pathway (KEGG database/hsa00010, gene numbers = 68) in patients with esophageal cancer (NCBI/GEO/GSE23400, n = 53). NES, normalized enrichment score. *P*-values were determined by Pearson’s correlation. **(E)** HEK293T cells were transfected with LOXL2-HA or L2Δ13-HA and aldolase A-Flag, GAPDH-Flag, enolase-Flag or empty vectors (Flag-vector and HA-vector). Whole cell lysates were co-immunoprecipitated with anti-Flag and probed by western blotting with the indicated antibodies. **(F)** GST pull-down assay in which either GST-tagged aldolase A or control GST was used to pull down endogenous LOXL2 in whole cell lysates extracted from KYSE510 cells. **(G)** Purified recombinant GST-aldolase A and GST from bacteria were retained on glutathione resins, incubated with either immunopurified LOXL2-Flag or L2Δ13-Flag from HEK293T cells and then immunoblotted with the antibody against Flag. **(H)** *In situ* PLA detection of the interaction between LOXL2 and aldolase A, L2Δ13 and aldolase A, and GAPDH with aldolase A in KYSE510 cells that express these proteins endogenously. PLA signals are shown in green, and nuclei are stained blue by DAPI dye. The detection of GAPDH and aldolase A complexes serves as a positive control. The negative control was performed with only anti-aldolase A antibody. Scale bar, 20 μm. **(I)** Top: illustration of the C- and N-terminal domains (CD and ND, respectively) of full-length LOXL2 and L2Δ13; bottom: co-IP assay of aldolase A with different domains of full-length LOXL2 and L2Δ13.

Our previous liquid chromatography tandem mass spectrometry (LC-MS/MS) analysis of proteins that co-purify with LOXL2 and/or L2Δ13 in esophageal cancer cells identified several known metabolism-associated proteins as interacting partners that interact physically with LOXL2/L2Δ13, including glycolic enzymes (16). Co-immunoprecipitation assays validated the interaction of LOXL2 and L2Δ13 with several such enzymes, including aldolase A, glyceraldehyde-3-phosphate dehydrogenase (GAPDH) and enolase (Fig. 4E; Supplementary Fig. S4C). Moreover, increased expression of these glycolic enzymes correlated with poor clinical outcome in esophageal cancer patients (Supplementary Fig. S4D and S4E). Importantly, pull-down, *in situ* proximity ligation assay (PLA) and immunofluorescence co-localization assays further validated that LOXL2 and L2Δ13 interact directly with aldolase A both *in vivo* and *in vitro*, suggesting independently that the classical enzyme activity of LOXL2 may not be required for its metabolic effects (Fig. 4F-4H; Supplementary Fig. S5A and S5B). To characterize the domains required for the binding of LOXL2/L2Δ13 to aldolase, we expressed in HEK293T cells a deletion mutant containing the shared N-terminal domain regions of LOXL2 and L2Δ13 (1-544 aa) as well as constructs expressing the C-terminus of LOXL2 (amino-acids 545-774) and L2Δ13 (amino-acids 545-729). The interaction of either LOXL2 or L2Δ13 with aldolase was largely abolished in HEK293T cells expressing each of the deletion mutants of LOXL2 and L2Δ13, suggesting that both the N- and C-terminal domains are required for efficient binding of LOXL2 and L2Δ13 to aldolase (Fig. 4I; Supplementary Fig. S5C). These findings indicate that LOXL2 and L2Δ13 may functionally drive tumor progression through induction of glycolysis as a result of interactions with glycolic proteins.

### LOXL2 and L2Δ13 promote the mobilization and enzymatic activity of aldolase A

To elucidate the molecular mechanisms by which LOXL2 and L2Δ13 modulate glycolysis, we focused on the glycolic enzymes that we have identified as enzymes associated with LOXL2/L2Δ13. Overexpression of either LOXL2 or L2Δ13 did not significantly alter the total protein expression levels of aldolase A, GAPDH and enolase *in vitro* and *in vivo* (Supplementary Fig. S6A and S6S6B). Aldolase A maintains glucose homeostasis depending on its enzymatic activity and contributes to glycolysis (31). Total aldolase activity was increased in nonmalignant esophageal epithelial cells expressing either LOXL2 or L2Δ13, as well as in L2Δ13-overexpressing mice (Supplementary Fig. S6C). In agreement with these observations, the catalytic activity of aldolase was inhibited following depletion of LOXL2 in esophageal cancer cells (Fig. 5A; Supplementary Fig. S6D). The decrease in glycolysis due to suppressed aldolase activity was fully reversed following re-expression of LOXL2/L2Δ13, raising the possibility that LOXL2 and L2Δ13 boost glycolysis by regulating aldolase activity (Fig. 5B).

**Figure 5.**
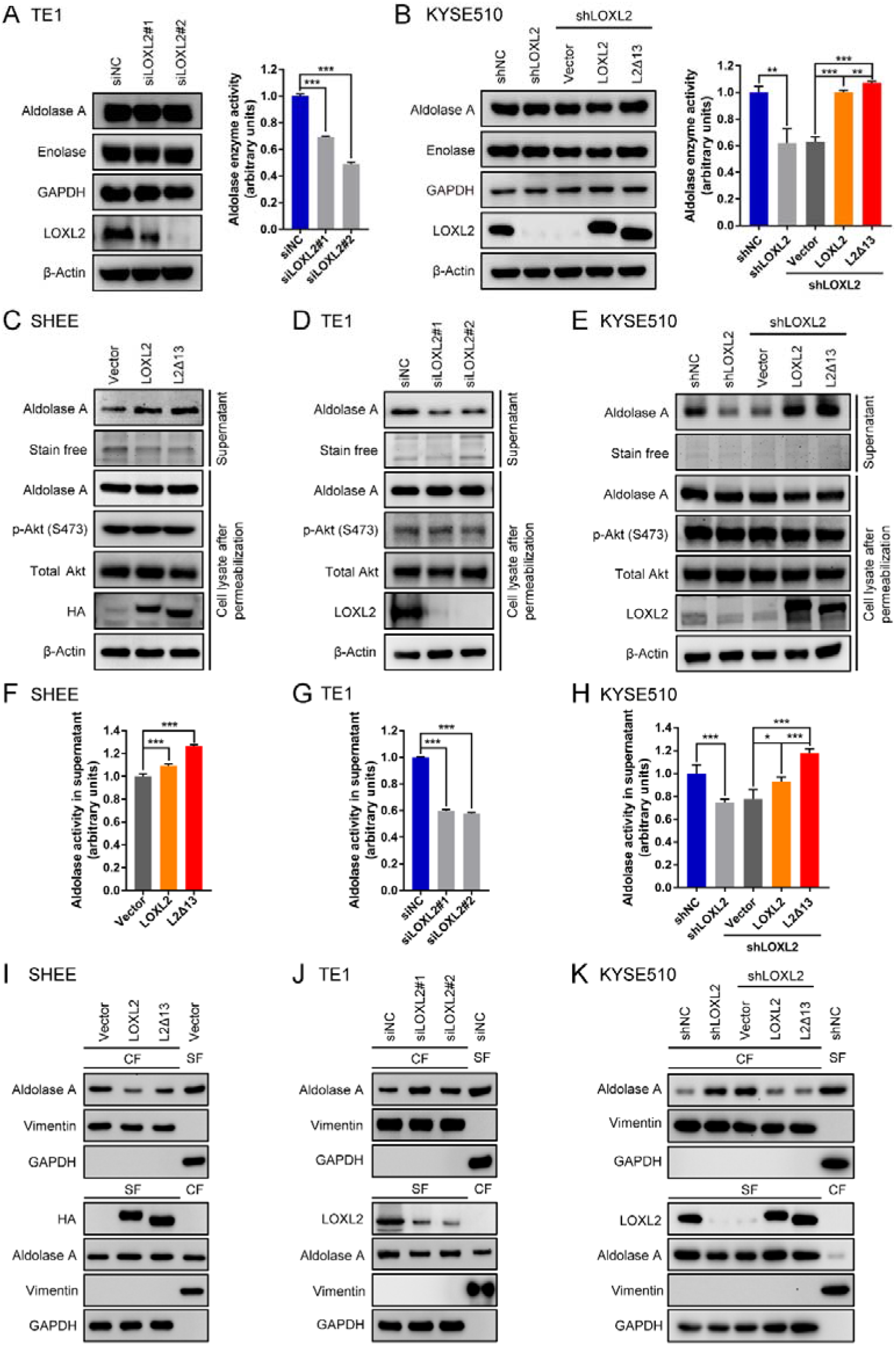
LOXL2 and L2Δ13 promote mobilization of aldolase A and increase its catalytic activity. **(A and B)** Western blotting detection (left) and aldolase enzyme activity assay (right) of esophageal cancer TE1 and KYSE510 cells upon LOXL2 silencing or re-expression of either LOXL2 or L2Δ13. **(C-E)** Nonmalignant esophageal epithelial SHEE cells and esophageal cancer cells (TE1 and KYSE510) were permeabilized with digitonin (30 μg/mL) for 5 min. Supernatant (top two panels) and cell lysate (bottom five panels) for each assay were subjected to western blotting as indicated. **(F-H)** Quantification of aldolase activity in the diffusible fraction (supernatant) by immunoblotting of cells in **(C-E)**. Bar graphs represent means ± SD, n = 3 or 4. \**P* < 0.05, \*\**P* < 0.01 or \*\*\**P* < 0.001 by *t*-test. **(I-K)** SHEE, TE1 and KYSE510 cells were lysed and fractionated into cytoskeletal fraction (CF) and soluble fraction (SF). Fractions from indicated cells transfected with empty vector, scrambled siRNA or shRNA are controls for the fractionation procedure. Vimentin served as a marker for the CF and GAPDH for the SF.

Glycolysis is organized around the actin cytoskeleton and binding of aldolase to filamentous actin inhibits its catalytic activity (22,32,33). To determine whether LOXL2/L2Δ13 interferes with the binding of aldolase A to the cytoskeleton, cells were permeabilized with digitonin to enable access of diffusible aldolase, followed by the collection of supernatant and cell lysates for further analyses of soluble and immobilized aldolase. Both LOXL2 and L2Δ13 induced the release of aldolase A in normal esophageal epithelial cells overexpressing cDNA encoding LOXL2 or L2Δ13 (Fig. 5C). Mobilization of aldolase A was prevented following the silencing of LOXL2 expression in esophageal cancer cells, which were conversely restored by re-expression of LOXL2 or L2Δ13 in cells silenced for LOXL2 expression (Fig. 5D and 5E; Supplementary Fig. S6E). The observed effects of LOXL2/L2Δ13 on glycolysis acceleration display similarities with the conventional effects of the serine/threonine kinase Akt on glycolysis (34). However, LOXL2/L2Δ13 mobilized aldolase A in an Akt-independent manner as illustrated by Akt S473 phosphorylation which remained constant regardless of the presence or absence of LOXL2 or L2Δ13 (Fig. 5C-5E; Supplementary Fig. S6E). Likewise, LOXL2- or L2Δ13-induced changes in the concentration of diffusible aldolase A were accompanied with a parallel alteration in the enzymatic activity of aldolase A in the supernatant derived from both nonmalignant and malignant cells (Fig. 5F-5H; Supplementary Fig. S6F). Cell fractionation indicated that both LOXL2 and L2Δ13 cause a shift of aldolase A from the cytoskeletal to the cytosolic fraction, whereas silencing LOXL2 expression inhibited the mobilization of aldolase A of esophageal cancer cells (Fig. 5I-5K; Supplementary Fig. S6G). Therefore, LOXL2/L2Δ13 stimulates aldolase mobilization to enhance its enzymatic activity, which in turn promotes glycolysis and tumor cell proliferation.

### LOXL2 and L2Δ13 deacetylate metabolic proteins including aldolase A

The acetylation level of proteins in KYSE510-derived cell lysates increased upon depletion of LOXL2, and was reduced following LOXL2/L2Δ13 re-expression (Fig. 6A; Supplementary Fig. S7A). A decay of global acetylation was also observed in L2Δ13-overexpressing mice as compared with wild-type mice (Supplementary Fig. S7B), suggestive of an extensive protein deacetylation function of LOXL2/L2Δ13. To monitor quantitative changes in LOXL2-regulated acetylation, we performed stable isotope labeling of amino acids in cell culture (SILAC) followed by affinity enrichment of lysine acetylation using nano-LC-MS/MS to systematically compare the acetyl peptide levels of KYSE510 cells before and after LOXL2 depletion (Fig. 6A). Proteomic analysis demonstrated that silencing LOXL2 expression altered the acetylation state of 249 proteins (331 sites) by greater than 1.5-fold, in which 241 (323 sites) were upregulated and only eight were downregulated (Supplementary Table S2). Proteins acetylated by LOXL2 depletion were mainly involved in metabolic processes including glucose metabolism, fatty acid metabolism and amino acid metabolism (Fig. 6B). Co-immunoprecipitation experiments were subsequently performed to validate the enhanced acetylation levels of some of the metabolic proteins that were identified in the proteomic screen. Intriguingly, only one of the LOXL2/L2Δ13-interacting proteins, aldolase A, was hyperacetylated in LOXL2-silenced esophageal cancer cells (Fig. 6C), while re-expression of LOXL2 or L2Δ13 reduced the elevated acetylation level of aldolase A (Fig. 6D). These results strongly suggest that LOXL2 and L2Δ13 induce aldolase deacetylation due to their physical interaction with aldolase A.

**Figure 6.**
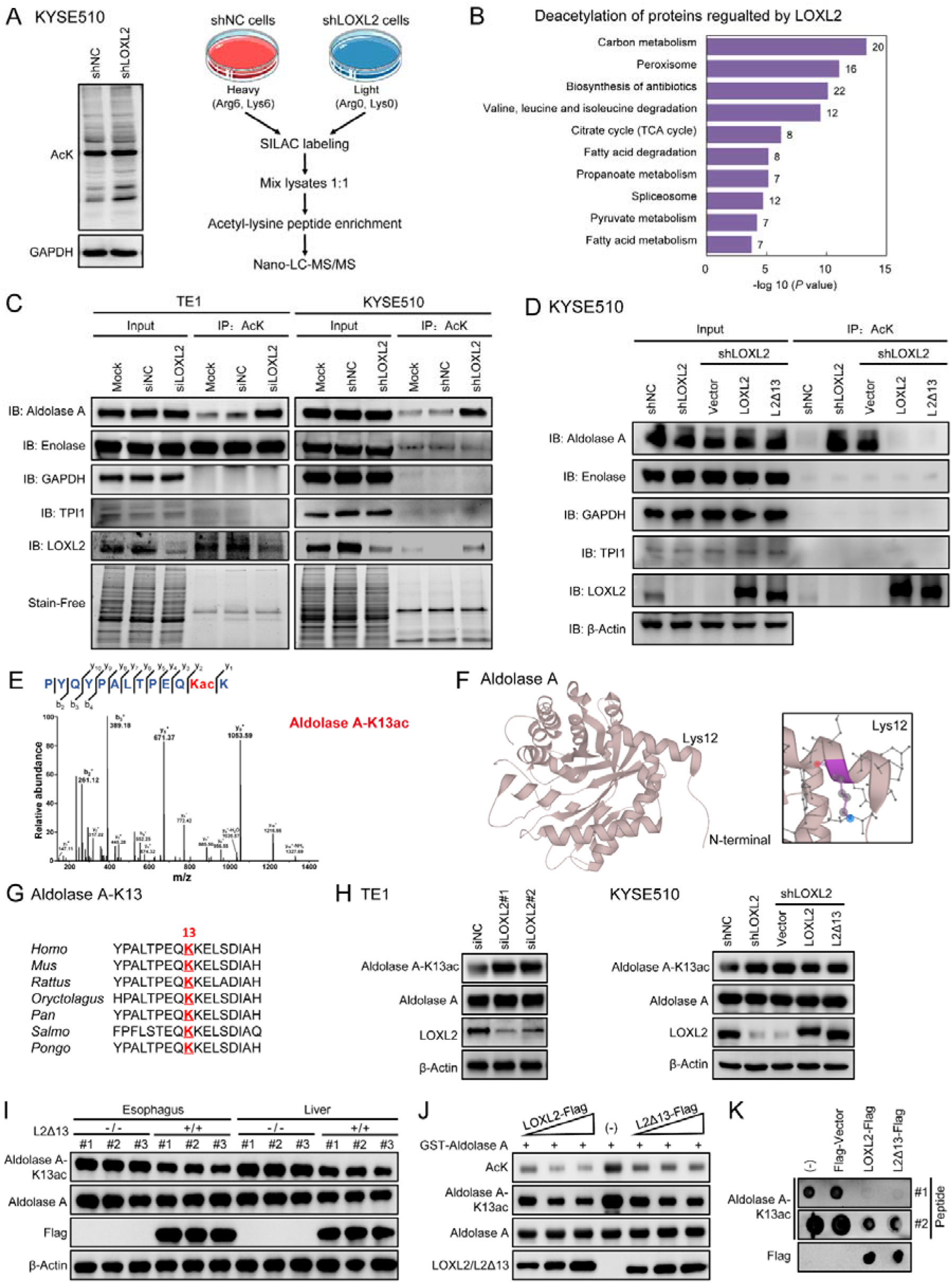
LOXL2/L2Δ13 catalyzes deacetylation of aldolase A-K13. **(A)** Left: acetylation level of total proteins from whole cell lysates of KYSE510 cells depleted for LOXL2; right: schematic diagram of the analytical strategy for SILAC labeling and global profiling of lysine acetylation. Control and LOXL2-depleted KYSE510 cells were separately labeled with “heavy” and “light” arginine and lysine using SILAC, and then proteins were digested for LC-MS/MS analysis. **(B)** Significant patterns show KEGG enrichment pathways of acetylated proteins in MS analysis. **(C)** Total proteins from untreated cells (mock), cells with scrambled siRNA or shRNA (siNC or shNC) and LOXL2-silenced cells were immunoprecipitated using acetyl-Lys and probed with indicated antibodies. Stain-Free gels was used as the control for equal protein concentration for the IP. **(D)** Total proteins from KYSE510 stably-infected cells were immunoprecipitated using acetyl-Lys and probed with indicated antibodies. Beta-Actin served as the control for equal protein concentration for the IP. **(E)** Endogenous aldolase A was purified from LOXL2-depleted and control KYSE510 cells with SILAC labeling, and acetyl peptides regulated by LOXL2 were identified. Shown are representative MS/MS spectra of acetylated aldolase A-K13. **(F)** Crystal structure model of human aldolase A (PDB database accession: 1ALD). Lys 12 in the model indicates ALDOA-K13 in our study. **(G)** Sequence alignment of acetylation sites of aldolase A-K13 from different species. **(H, I)** Aldolase A acetylated at K13 (aldolase A-K13ac) and total aldolase A expression level in indicated cells and esophagi and livers of mice. **(J)** Purified GST-aldolase A from bacteria were incubated with increasing amount of purified LOXL2-Flag or L2Δ13-Flag from HEK293T transfectants for *in vitro* LOXL2/L2Δ13 deacetylase activity assays. **(K)** Indicated acetyl peptides of the aldolase A-K13 were incubated with the purified LOXL2-Flag, L2Δ13-Flag and Flag-vector proteins from HEK293T transfected cells. The reaction products were blotted with antialdolase A-K13ac or anti-Flag.

To identify the lysine residues of aldolase A that are deacetylated by LOXL2/L2Δ13, we purified SILAC labeled endogenous aldolase A from LOXL2-depleted and control KYSE510 cells. The K13 residue of aldolase A was hyperacetylated in LOXL2-depleted cells but not in control cells (Fig. 6E; Supplementary Table S2). According to sequence alignment analysis, the K13 residue (K12 in the PDB-1ALD structure model) is highly conserved between diverse animal species (Fig. 6F and 6G). We further synthesized acetyl peptides derived from the aldolase A-K13 and generated polyclonal rabbit antibodies specifically directed against the acetylated K13 residue of human aldolase A (Supplementary Fig. S7C). Aldolase A-K13 was frequently acetylated in nonmalignant cells and different types of esophageal cancer cells (Supplementary Fig. S7D). Interestingly, depletion of LOXL2 prominently enhanced the acylation of aldolase-K13 (Fig. 6H; Supplementary Fig. S7E). Overexpression of LOXL2 or L2Δ13 conversely inhibited aldolase A-K13 acetylation in both cancer cells and mice (Fig. 6H and 6I; Supplementary Fig. S7E). To directly measure the aldolase A-targeted deacetylase activity of LOXL2 and L2Δ13, we performed an *in vitro* aldolase deacetylation assay using purified Flag-tagged LOXL2 or Flag-tagged L2Δ13 and GST-tagged aldolase A proteins. Both LOXL2 and L2Δ13 efficiently catalyzed aldolase A deacetylation at the K13 locus in a concentration-dependent manner (Fig. 6J; Supplementary Fig. S7F). In agreement with these observations, two different K13 acetyl peptides were also specifically deacetylated by LOXL2 and L2Δ13 *in vitro* (Fig. 6K). These results identify aldolase A as a LOXL2 deacetylation substrate and demonstrate that this activity of LOXL2 is independent from its conventional lysyloxidase activity.

### LOXL2/L2Δ13-dependent deacetylation of aldolase A-K13 induces metabolic reprogramming in esophageal cancer progression

The acetylation levels of glycolic enzymes correlate with their enzymatic activities in primed human pluripotent stem cells (35). Neither the acetyl-mimetic mutant of aldolase, K13Q, nor its non-acetylatable mutant, K13R, had any direct effect on the aldolase enzymatic activity in esophageal cancer cells (Fig. 7A). However, the K13Q acetyl-mimetic mutant, but not the K13R non-acetylatable mutant, significantly prevented the release of aldolase A in the cytoplasm (Fig. 7B). Cell fractionation experiments confirmed that aldolase acetylation at K13 impeded the dissociation of aldolase A from the cytoskeletal compartment, which was further verified by colocalization analysis of aldolase A and actin fibers (Fig. 7C and 7D). These observations suggest that deacetylation of aldolase A by LOXL2 and L2Δ13 promotes the mobilization of aldolase from the actin cytoskeleton, thus aldolase A-K13 deacetylation plays an important role in the regulation of glycolysis.

**Figure 7.**
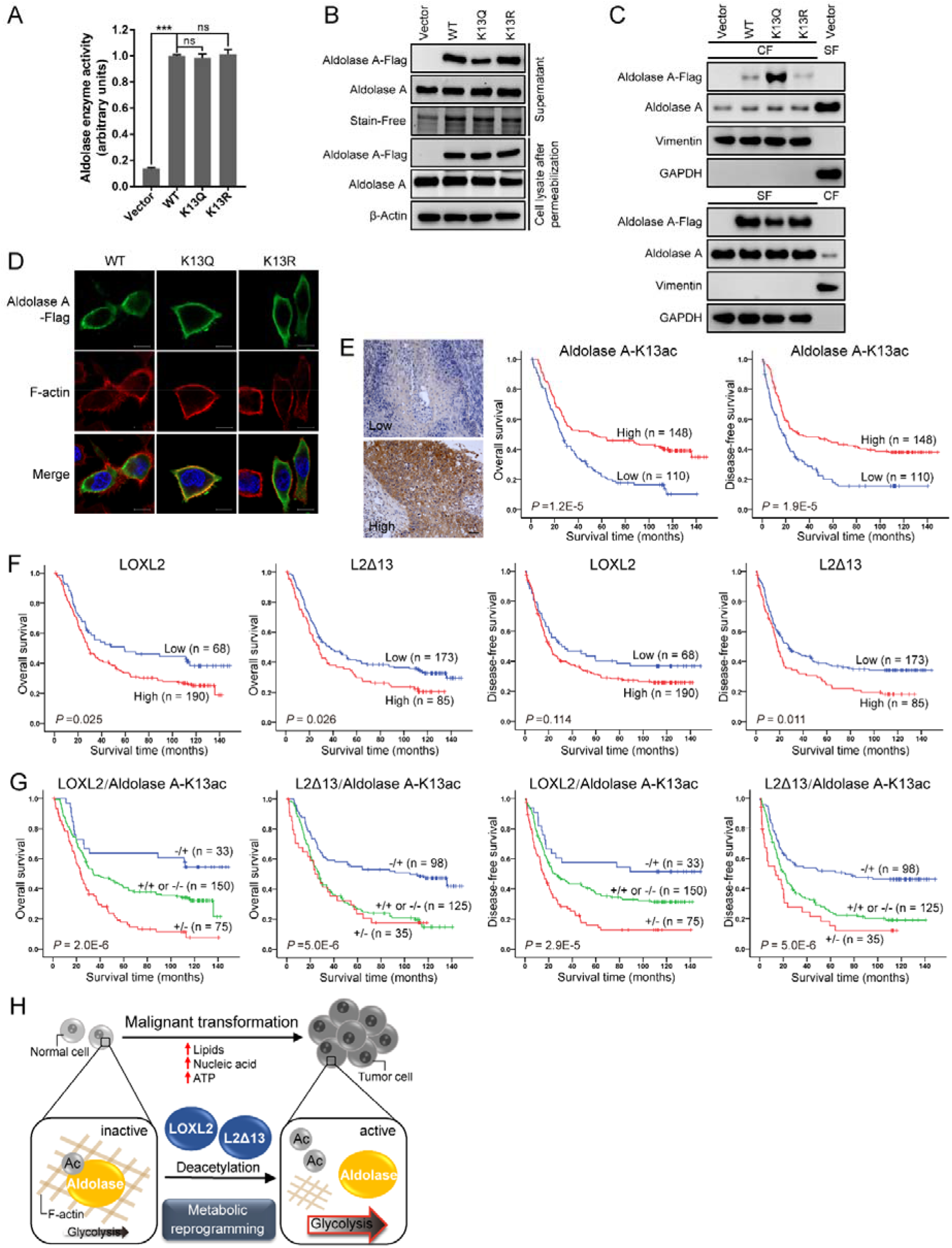
LOXL2-dependent deacetylation of aldolase A-K13 induces metabolic reprogramming in esophageal cancer progression. **(A)** Flag-tagged wild-type aldolase A, acetyl-mimetic mutant aldolase A-K13Q and non-acetylatable mutant aldolase A-K13R were expressed in KYSE510 cells. Aldolase A proteins were purified by IP with Flag antibody, and aldolase activity was determined. Cells transfected with Flag-tagged vector served as the negative control. Data represent mean ± SD (n = 3). \*\*\**P* < 0.001; ns, not significant. **(B)** Western blotting of supernatant and cell lysate of KYSE510 cells transfected with empty vector, Flag-tagged aldolase A (WT), its acetyl-mimetic or non-acetylatable mutant, including K13Q and K13R. **(C)** KYSE510 cells expressing Flag-tagged wild-type aldolase A, its acetyl-mimetic mutant K13Q and non-acetylatable mutant K13R were lysed and fractionated. Vimentin is used as a marker for the CF and GAPDH for the SF. Fractions from the cells transfected with Flag-tagged empty vector are controls for the fractionation procedure. **(D)** Immunofluorescence analysis of the effect of different aldolase A mutants from **(B)** on its release from the actin cytoskeleton in KYSE510 cells. Scale bar, 10 μm. **(E)** Representative low and high acetylation of aldolase A-K13 by IHC analysis (left) and Kaplan-Meier curves by log-rank test (right) of aldolase A-K13ac in tissue microarrays of patients with esophageal cancer (n = 258). Scale bar, 50 mm. **(F)** Kaplan-Meier curves by log-rank test of LOXL2 and L2Δ13 for overall survival and disease-free survival in patients of **(E)**. **(G)**Kaplan-Meier estimates of overall survival and disease-free survival of patients with esophageal cancer according to LOXL2 or L2Δ13 associated with acetylated aldolase A-K13. **(H)** Proposed model integrating the LOXL2/L2Δ13-activated metabolic reprogramming through deacetylating glycolic proteins in tumorigenesis and tumor progression.

The contribution of the acetylation status of aldolase-K13 to glycolysis in esophageal cancer cells prompted us to determine its prognostic value in clinical settings. We therefore measured the levels of K13 acetylated aldolase A by immunohistochemical staining of tumor sections derived from 258 esophageal cancer patients. Patients displaying high level acetylation of aldolase A had longer overall survival and disease-free survival as compared with patients in which aldolase A was less acetylated (Fig. 7E). Moreover, hyperacetylated aldolase A correlated with histologic grade and tumor invasion state and was found to represent an independent prognostic factor for favorable clinical outcome (Supplementary Table S3 and S4). Consistently with our previous study (16), elevated LOXL2 or L2Δ13 was significantly associated with poor survival in these patients (Fig. 7F). We subsequently combined the states of either LOXL2 or L2Δ13 and K13 acetylated aldolase A to develop molecular models for predicting the clinical prognosis of patients with esophageal cancer. Indeed, low level expression of LOXL2 or L2Δ13 was associated with increased acetylation of aldolase A-K13, and was significantly correlated with longer overall survival and disease-free survival (Fig. 7G).

To conclude, our experiments identify LOXL2 as a novel deacetylase that directly interacts and deacetylates aldolase A. This in turn induces its release from filamentous actin and enzymatic activity, leading to accelerated glycolysis which subsequently contributes to tumor development and tumor metastasis (Fig. 7H).

## Discussion

Previous studies have shown that the pro-fibrogenic effects of LOXL2 are dependent on its extracellular amine oxidase activity. On the other hand, the pro-tumorigenic effects of LOXL2 are mediated by a combination of extracellular and intracellular activities of LOXL2 (3,4,7–10). Here we generated a genetically engineered gain-of-function mouse model for *L2Δ13*, a non-secreted catalytically inactive splice form of *LOXL2,* and found that L2Δ13 is able to regulate metabolic reprogramming so as to promote tumor initiation and progression *in vitro* and *in vivo* despite its lack of amine oxidase activity. We further showed that both L2Δ13 and LOXL2 bind to several glycolysis-promoting enzymes, including aldolase A. In the case of aldolase A, this interaction triggered the deacetylation of aldolase on the K13 residue. This in turn led to enhanced glycolysis that ultimately contributes to tumor growth and tumor metastasis in esophageal cancer.

Germ-line overexpression of full-length LOXL2 causes male sterility due to testicular degeneration and epididymis dysfunction (10). However, the offspring of homozygous L2Δ13 overexpressing mice do not display such reproductive dysfunction and conform to the expected Mendelian ratio, indicating that full-length LOXL2-induced male sterility is probably linked to its lysyl oxidase activity. L2Δ13-overexpressing mice exhibited reduced body weight, which was accompanied by the disruption of glucose and lipid homeostasis. This observation was further validated by the integrated transcriptomic and metabolomic analysis of L2Δ13-induced metabolic pathway changes in transgenic mouse models. Full-length LOXL2 is known to promote cardiac, liver and renal fibrosis (3,4,36). We demonstrated here that the pro-fibrotic role of LOXL2 is partially due to its novel deacetylase activity rather than to its classical lysyl-oxidase activity. Compared with wild-type mice, L2Δ13-overexpressing mice exhibit higher sensitivity to carcinogens such as CCl4 and NMBA, resulting in a higher risk of tumorigenic transformation. These results provide strong evidence for the physiological and pathological relevance of the amine oxidase activity independent effects of LOXL2 and its enzymatically inactive splice forms.

Tumor cells preferentially uptake, transfer and utilize glucose at much higher rates in order to generate ATP, maintain redox balance and support biosynthesis, thereby reprogramming their metabolism and that of other cells in the tumor microenvironment, as compared with the metabolism of normal cells (37). Loss- and gain-of-function assays showed that, in addition to the effects of LOXL2 on tumor cell invasion and tumor metastasis, both L2Δ13 and LOXL2 promote esophageal cancer cell proliferation and tumor growth in xenograft experiments (16,38–40). The Warburg effect is characterized by enhanced glycolysis and lactate production regardless of oxygen availability, and represents a hallmark of cancer cells (30,41). Interestingly, we found that a similar enhancement of glycolysis takes place in normal esophageal epithelial cells following overexpression of either LOXL2 or L2Δ13. In contrast, depletion of LOXL2 in esophageal cancer cells inhibits glycolysis, suggesting that glycolysis induced by L2Δ13/LOXL2 contributes to the Warburg effect and tumor progression.

We identified and validated aldolase A, GAPDH, and alpha-enolase as glycolysis-associated proteins that physically interact with LOXL2 and L2Δ13. Aldolase A catalyzes the conversion of fructose-1,6-bisphosphate (FBP) to glyceraldehyde-3-phosphate (GA3P) and dihydroxyacetone phosphate (DHAP), providing a major source of substrates required for nucleotide and purine biosynthesis (31). Our findings suggest that both the accelerated glycolysis and cell proliferation observed in esophageal cancer is due, at least in part, to the interactions of LOXL2/L2Δ13 with aldolase A. Aberrant expression of aldolase A is found to be associated with tumor metastasis and worse patient prognosis in several types of solid tumors, including lung cancer and pancreatic ductal adenocarcinoma (42,43). Inhibition of aldolase A is sufficient to block glycolysis, thereby inhibiting the uncontrolled cell proliferation of tumor cells under hypoxic conditions (44). We found that silencing L2Δ13/LOXL2 expression inhibits the enzymatic activity of aldolase A without affecting its expression. Aldolase A binds to and accumulates on actin fibers through electrostatic bonds associated with its FBP substrate (22,32,33). Our results suggest that L2Δ13 and LOXL2 control the release of aldolase from filamentous actin and simultaneously stimulate the enzymatic activity of aldolase A via its deacetylation on K13. We have previously demonstrated that cytoplasmically localized LOXL2, as well as L2Δ13, induces cytoskeletal reorganization to promote tumor invasion and metastasis (16). This LOXL2/L2Δ13-induced reorganization may also trigger aldolase release into the cytoplasm to further enhance tumor progression. Taken together, our results strongly demonstrate that the enzyme-independent pro-tumorigenic effects of L2Δ 13/’LOXL2 on glycolysis and cytoskeletal dynamics contribute to tumor progression.

Evolutionarily conserved lysine acetylation regulates the stability, activity, and localization of metabolic enzymes in response to extracellular changes (45). Our results concerning the deacetylase activity of LOXL2 are in agreement with prior studies on the contributions of the deacetylase activity of LOXL3 to inflammatory responses (46). Classical deacetylases, such as the histone deacetylases (HDACs) and sirtuins (SIRTs), have recently been found to either directly or indirectly modify the acetylation state of lysine residues associated with some glycolytic enzymes (47). L2Δ13/LOXL2 deacetylates aldolase on K13 and facilitates its release from the filamentous actin, which may be due to the location of K13 in a positively-charged surface region of aldolase that mediates its attachment to actin (33).

To summarize, we provide compelling evidence that identifies LOXL2 and L2Δ13 as novel deacetylates with physiological and pro-tumorigenic functions in cell metabolism. Our findings highlight a hitherto-unknown mechanism by which LOXL2 and L2Δ13 catalyze the deacetylation of aldolase to stimulate its mobilization and enzymatic activity in glycolysis that in turn promotes tumor progression.

## Acknowledgments

We are very thankful to Professor Stanley Li Lin of Shantou University Medical College for editing the manuscript. This work was supported by grants from the National Natural Science Foundation of China (81472613 and 81872372 to E.-M. Li; 81772532 to L.-Y. Xu; 82103495 to X.-H Zhan), Li Ka Shing Foundation Grant for Joint Research Program between Shantou University and Technion-Israel Institute of Technology (43209506 to G. Neufeld and E.-M. Li), Special Fund Project for Science and Technology Innovation Strategy of Guangdong Province (210713096870791 to X.-H Zhan), Li Ka Shing Foundation Cross-Disciplinary Research Grant (2020LKSFG07B to E.-M. Li), the National Cohort of Esophageal Cancer of China (2016YFC0901400 to L.-Y. Xu) and the National Key R&D Program of China (2018YFC1313100 to E.-M. Li).

## Author Contributions

J.-W. Jiao and X.-H. Zhan designed and performed the majority of the experiments, analyzed and interpreted the data, and wrote the manuscript. J.J Wang and L.-X He performed some mouse experiments and pathology analysis. Z.-C. Guo performed proteomic analysis of cancer cells. X.-E. Xu performed H&E and immunohistochemistry experiments. L.-D Liao and B. Wen assisted with cell culture and immunofluorescence. X. Huang and Y.-W. Xu performed blood chemistry analysis. H. Hu and Z.-J. Chang provided critical advice and consultations. G. Neufeld revised the manuscript and provided expertise on oncology. K. Zhang, L.-Y. Xu and E.-M. Li designed the study, provided advice and reagents, and directed the study.

## Notes

The authors declare no potential conflicts of interest.

### Competing Interest Statement

The authors have declared no competing interest.

