## Supplementary figures and images for "LOXL2-dependent deacetylation of aldolase A induces metabolic reprogramming and tumor progression"

### Supplementary Figures

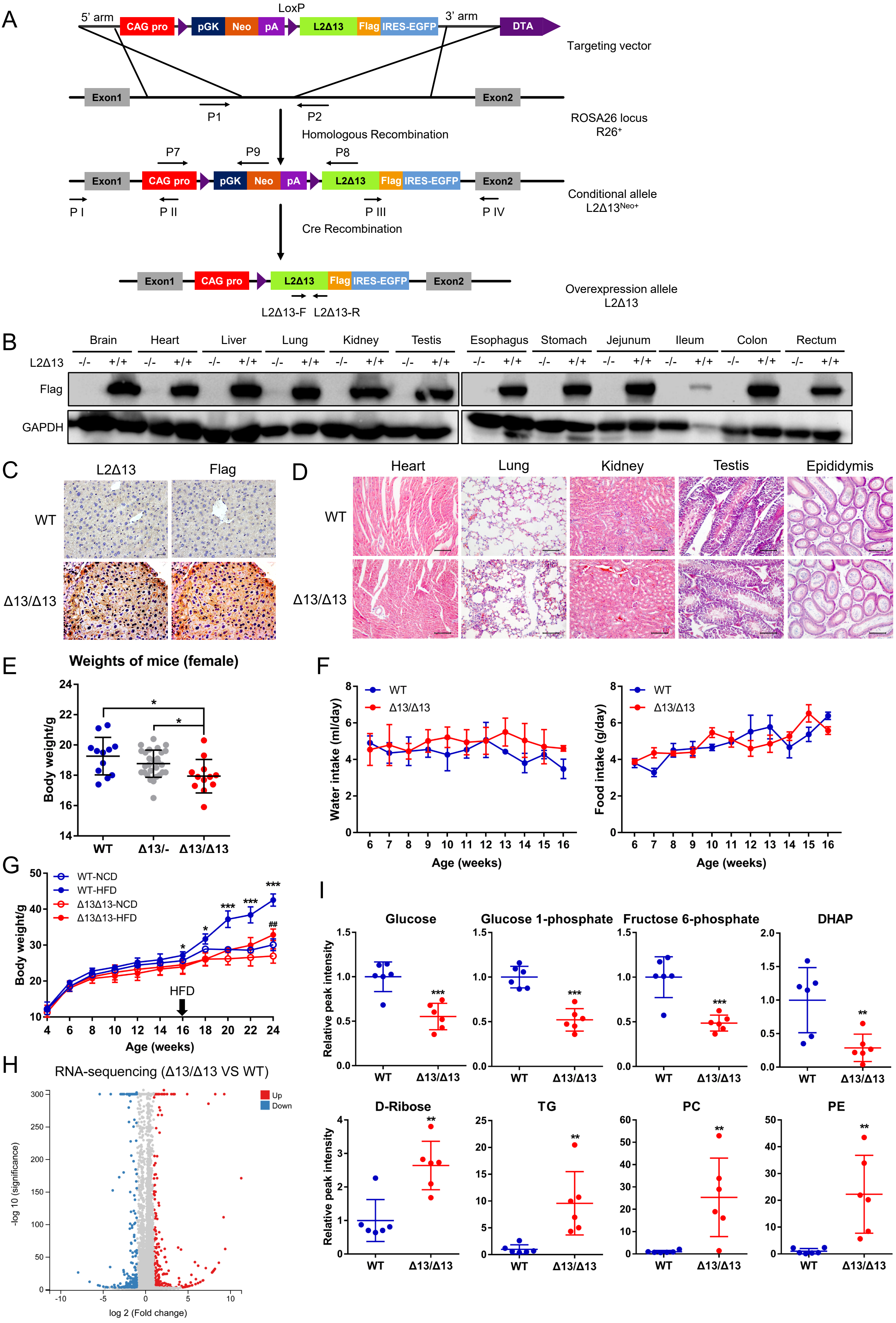

#### Figure S2

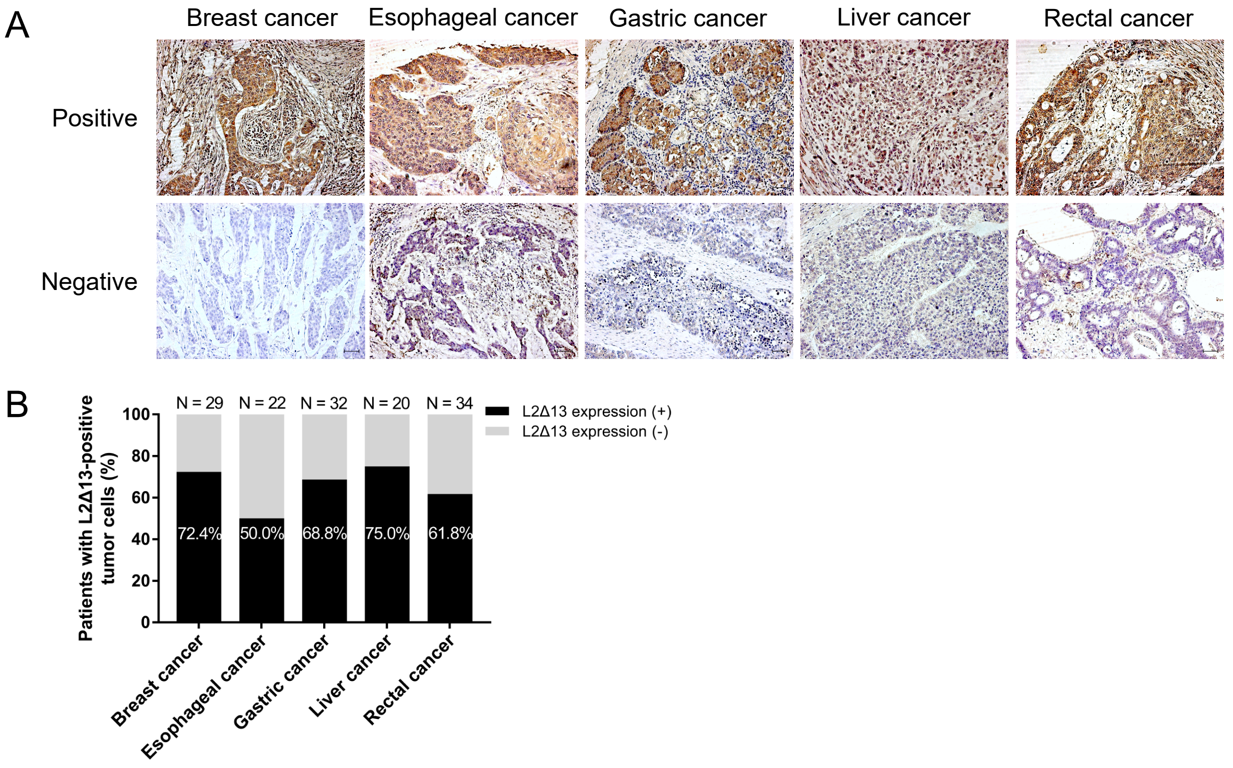


#### Figure S3

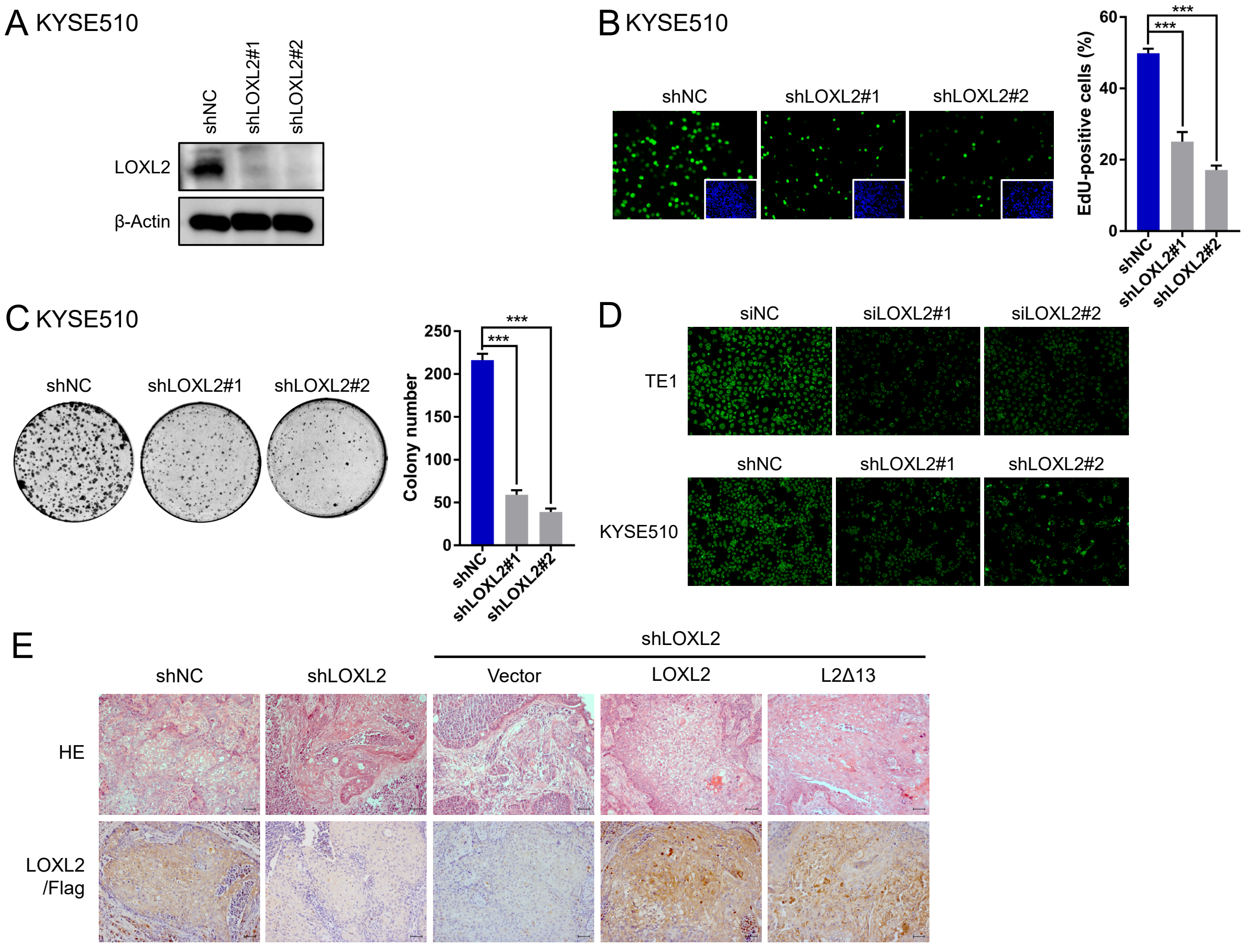

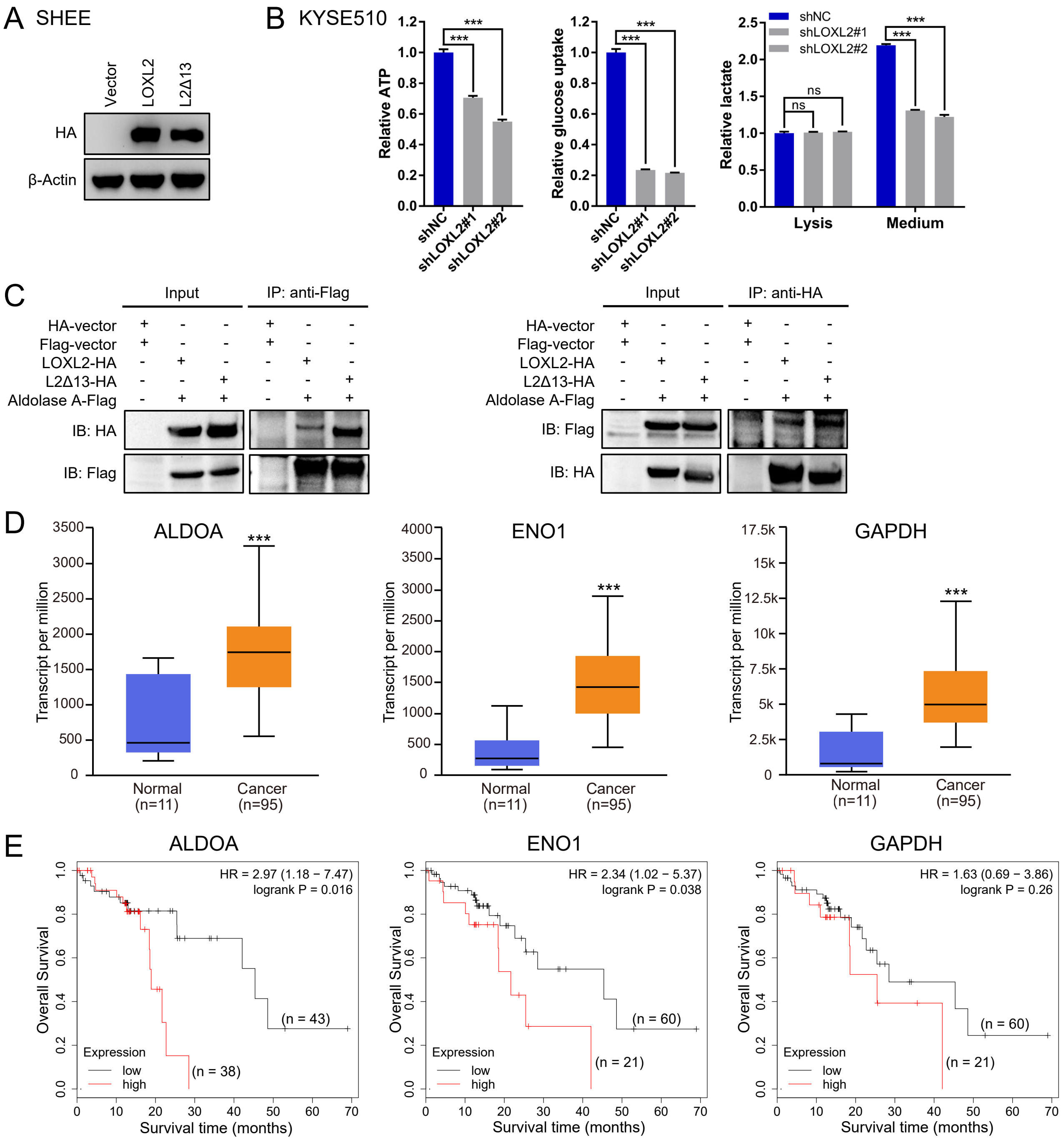

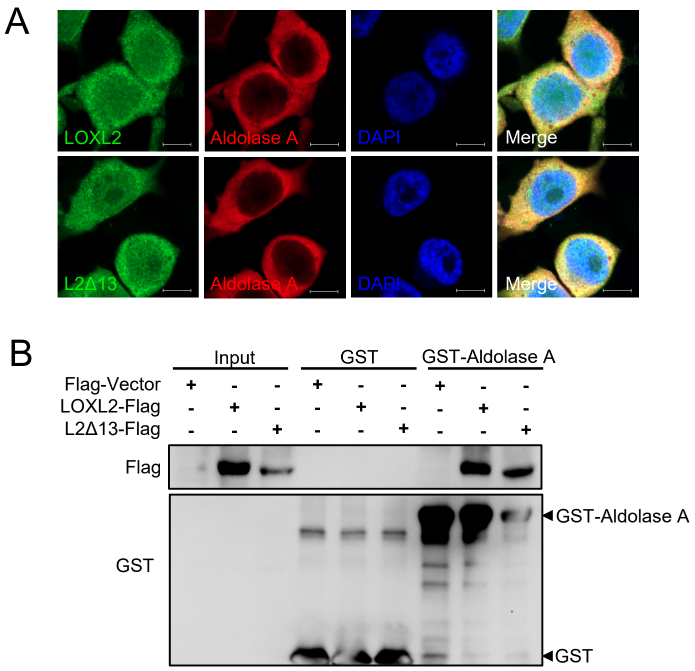

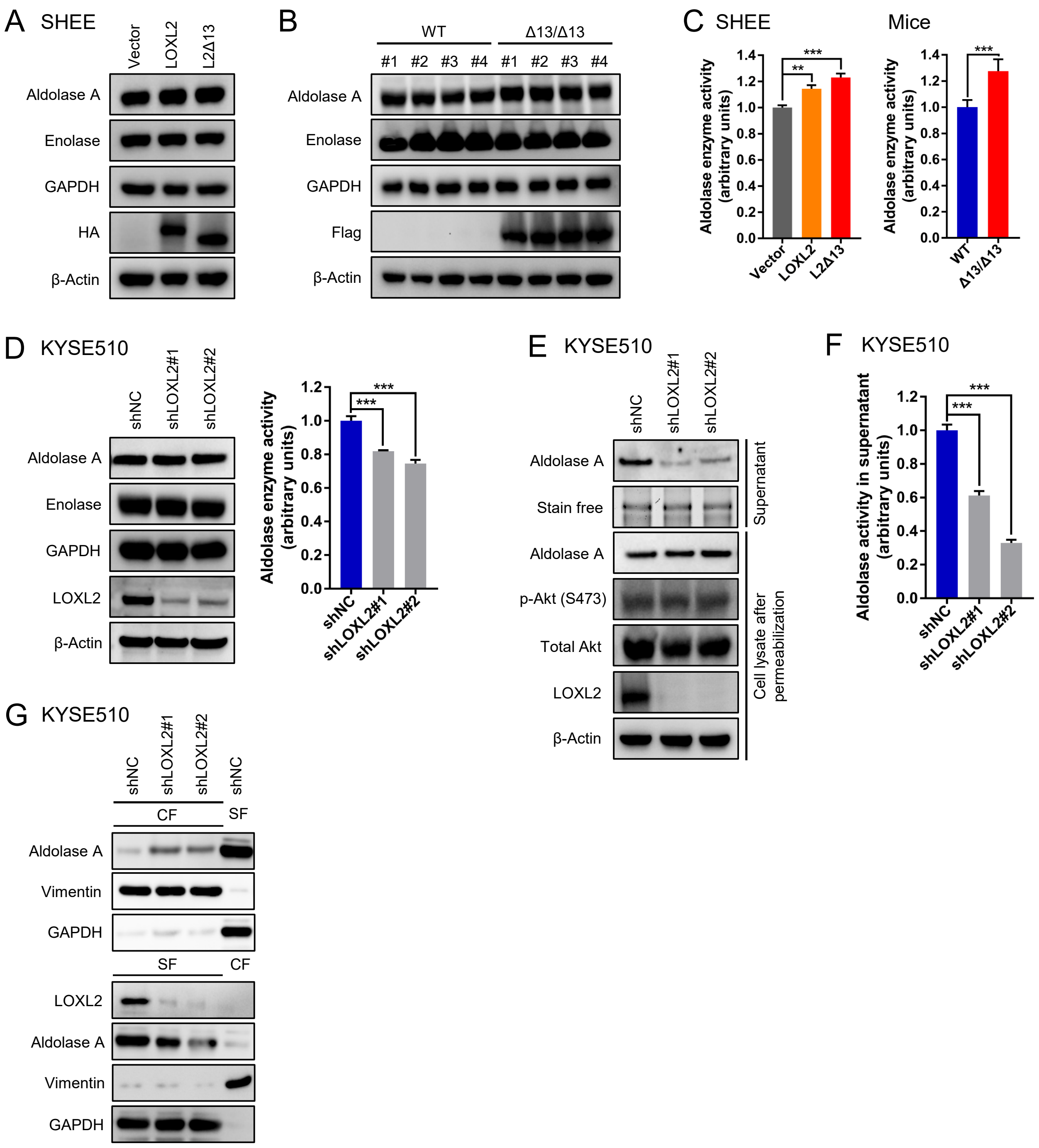

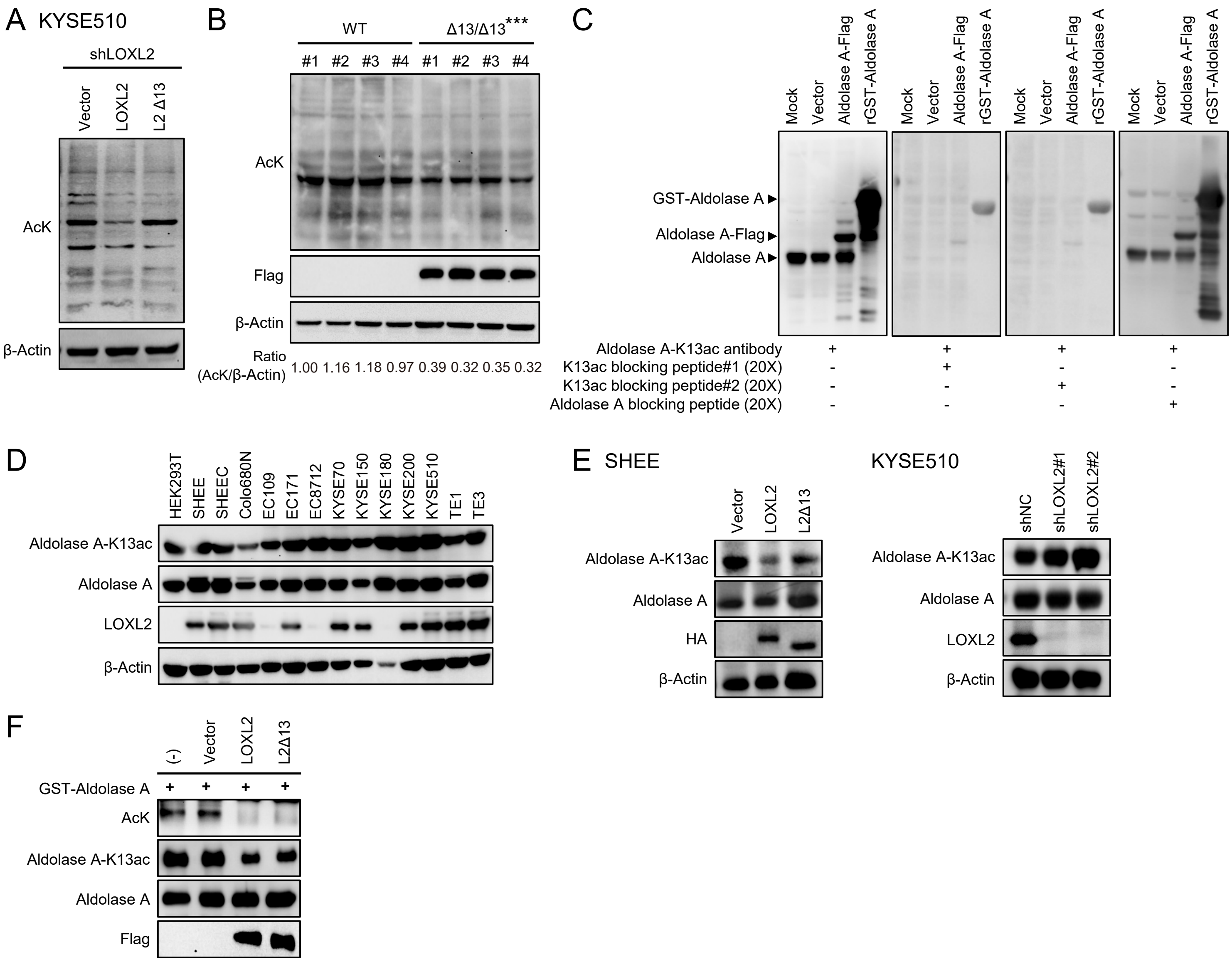
