## Supplementary Figure Legends for "LOXL2-dependent deacetylation of aldolase A induces metabolic reprogramming and tumor progression"

**Figure S1.** **Generation and comparison of L2Δ13-overexpressing transgenic mice and control wild-type mice.**

**(A)** Schematic diagram illustrating different *ROSA26* allele variants used in the study. The targeting vector contains a *PGK-neomycin-polyA* element flanked by *LoxP* sites, and the human *L2Δ13* cDNA sequence fuses to the 3X Flag sequence and the *IRES-EGFP* reporter gene. An *L2Δ13^Neo+^* allele is initially generated by homologous recombination. The *PGK-neomycin-polyA* is removed upon Cre-mediated excision, then the *L2Δ13/Flag* is expressed under the control of the CAG promoter (*L2Δ13* allele). Primer pairs for PCR analysis are listed in *Supplementary Table S5*. **(B)** Expression of L2Δ13-Flag from different tissues was assayed by western blotting in wild-type and homozygous L2Δ13-overexpressing mice. **(C)** Representative histochemical staining with antibodies against L2Δ13 or Flag for livers derived from the wild-type and L2Δ13-overexpressing mice. Scale bar, 50 μm. **(D)** Representative hematoxylin and eosin (H&E) staining of different organs from wild-type and L2Δ13-overexpressing mice, including heart, lung, kidney, testis and epididymis. Scale bar, 50 μm. **(E)** Body weight of female wild-type, heterozygous and homozygous L2Δ13-overexpressing mice at 8 weeks of age (n = 12, 28, 12). **P* < 0.05. **(F)** Water intake and food intake of wild-type and Δ13/Δ13 mice were monitored at the indicated time points (n = 8 per group). **(G)** Body mass of male Δ13/Δ13 mice and littermate wild-type controls fed either a normal chow diet (NCD) or high-fat diet (HFD) for 8 weeks (n = 3 or 4). *P*-values were determined using a *t*-test. ^*^Differences between two groups of wild-type mice, **P* < 0.05 or ****P* < 0.001. ^#^Differences between two groups of Δ13/Δ13 mice, ^##^*P* < 0.01. **(H)** Volcano plot of differential gene expression between L2Δ13-overexpressing and age-matched wild-type control mice in RNA-sequencing analysis (n = 6; GSE145238). **(I)** Representation of the downregulated glycolysis/gluconeogenesis pathways (top) and the upregulated pentose phosphate and fatty acid metabolism pathways (bottom) and relative MS peak intensity of their corresponding intermediate metabolites. ***P* < 0.01; ****P* < 0.001. *P*-value was calculated using a *t*-test. DHAP, dihydroxyacetone phosphate; TG, triglyceride; PC, phosphatidyl choline; PE phosphatidyl ethanolamine.

**Figure S2. The expression of L2Δ13 protein in patients with different types of cancer.**

**(A)** Representative paraffin sections stained with L2Δ13 antibody for patients with breast cancer, esophageal cancer, gastric cancer, liver cancer and rectal cancer. Scale bar, 50 μm. **(B)** Proportions of patients with L2Δ13-positive tumor cells in different cancers from **(A)**.

**Figure S3. L2Δ13 and LOXL2 promote tumor cell proliferation *in vitro* and *in vivo*.**

**(A-C)** Western blotting **(A)**, EdU **(B)** and colony formation **(C)** assays with esophageal cancer KYSE510 cells following LOXL2 knockdown with a specific shRNA target. Error bars indicate mean ± SD of three replicates. ****P* < 0.001 by *t*-test analysis. **(D)** Effects of LOXL2 silencing on lipid droplets in TE1 and KYSE510 cells. **(E)** Representative paraffin sections stained with hematoxylin and eosin (H&E, top) and antibodies against LOXL2 or Flag (bottom) for tumors derived from the xenografts. LOXL2 antibody was used for the section from xenografts implanted with KYSE510 cells expressing either a scrambled (shNC) or LOXL2-silencing lentiviral vector (shLOXL2), while the Flag antibody was adopted for the remaining three groups. Scale bar, 50 μm.

**Figure S4. LOXL2 and L2Δ13 drive glycolysis by interacting with glycolic proteins.**

**(A)** Western blotting assays of nonmalignant esophageal epithelial cells expressing the empty vector, full-length LOXL2 or L2Δ13 variant. **(B)** Levels of ATP, glucose uptake and lactate of esophageal cancer cells following LOXL2 silencing by specific shRNA. Data show the mean ± SD (n = 3 or 4). ****P* < 0.001 by the *t*-test. ns, not significant. **(C)** Co-IP assays of LOXL2-HA or L2Δ13-HA with aldolase A-Flag. Cell lysates were subjected to immunoprecipitation with anti-Flag (left) or anti-HA (right) and probed by western blotting with the indicated antibodies. **(D)** RNA-sequencing data of gene expressions of ALDOA, ENO1 and GAPDH from TCGA database in esophageal cancer (n = 95) and normal esophagus (n=11). ****P* < 0.001 by the *t*-test. **(E)** Kaplan-Meier estimates overall survival of patients with esophageal cancer according to curves of ALDOA, ENO1 and GAPDH (n = 81; Kaplan-Meier Plotter database, http://kmplot.com/analysis/). High expression levels of these glycolic genes were associated with shorter medium survival time in patients with esophageal cancer. Statistical significance was assessed by log-rank test. HR, hazard ratio (95% CI).

**Figure S5. LOXL2 and L2Δ13 directly interact with aldolase A.**

**(A)** Confocal immunofluorescence staining indicate co-localization of either LOXL2 or L2Δ13 with aldolase A in KYSE510 cells that express these proteins endogenously. Shown are individual views of LOXL2/L2Δ13 (green), aldolase A (red) and nucleus (blue), and merged images showing triple staining. Scale bar, 10 μm. **(B)** GST-aldolase A and GST were retained on glutathione resins, incubated with whole cell lysates extracted from HEK293T cells transfected with LOXL2-Flag, L2Δ13-Flag or Flag empty vector, and then subjected to western blotting as indicated. **(C)** Pull-down assay in which either GST-tagged aldolase A or control GST was used to pull down different HA-tagged deletion mutants of full-length LOXL2 and L2Δ13 in whole cell lysate from HEK293T cells expressing each of these mutants.

**Figure S6. Depletion of LOXL2 inhibits the mobilization and enzymatic activity of aldolase A.**

**(A)** Western blotting analysis of nonmalignant esophageal epithelial SHEE cells expressing HA-tagged LOXL2, HA-tagged L2Δ13 or the empty vector. **(B)** Immunoblotting detection of wild-type and homozygous L2Δ13-overexpressing mice (n = 4). **(C)** Aldolase enzyme activity analysis of SHEE cells overexpressing LOXL2 or L2Δ13 (left; n = 3), wild-type and L2Δ13-overexpressing mice (right; n = 4). Data show means ± SD, ****P* < 0.001. **(D)** Western blotting (left) and aldolase activity determination (right) of KYSE510 esophageal cancer cells upon depletion of LOXL2. **(E)** KYSE510 cells were permeabilized with digitonin (30 μg/mL) for 5 min. Supernatant and cell lysate were subjected to immunoblotting. **(F)** Quantification of aldolase activity in the supernatant by immunoblotting of cells from **(E)**. Means ± SD, n = 3. ****P* < 0.001 by the *t*-test. **(G)** KYSE510 cells following depletion of LOXL2 were lysed and fractionated. Vimentin served as a marker for the cytoskeletal fraction (CF) and GAPDH for the soluble fraction (SF). Fractions from the cells transfected with scrambled shRNA are controls for the fractionation procedure.

**Figure S7. LOXL2 and L2Δ13 catalyze deacetylation of aldolase A-K13.**

**(A)** Acetylation level of total proteins from whole cell lysates of stably LOXL2-silenced KYSE510 cells following LOXL2/L2Δ13 re-expression by Western blotting with anti-acetyl-Lys antibody. **(B)** Acetylation level of total proteins from livers of wild-type and L2Δ13-overexpressing mice. Ratios of AcK/β-actin between two groups were quantifiably analyzed by *t*-test, ****P* < 0.001. **(C)** Specificity of antibody directed against the acetylation of aldolase A-K13 was detected by Western blotting in untreated HEK293T cells, HEK293T cells expressing Flag-tagged aldolase A or Flag-tagged empty vector and purified recombinant GST-aldolase A from bacteria. Positive staining of aldolase A-K13ac was strongly blocked by indicated matching blocking peptides (#1 and #2) to the aldolase A-K13ac antibody, but not by the peptide to the total aldolase A antibody. **(D)** The expression levels of aldolase A-K13ac, total aldolase A and full-length LOXL2 in nonmalignant cells (HEK293T and SHEE) and different types of esophageal cancer cells. **(E)** Western blotting analysis of aldolase A acetylated at K13 (aldolase A-K13ac) and total aldolase A in SHEE cells expressing HA-tagged LOXL2 or HA-tagged L2Δ13 and KYSE510 cells silenced for LOXL2 expression. **(F)** Flag-tagged LOXL2, L2Δ13 and empty vector proteins were purified from HEK293T transfectants using Flag antibody, and then incubated with GST-aldolase A purified from bacteria in the LOXL2/L2Δ13 reaction buffer for in vitro deacetylase activity assay.
