## Supplementary Materials and Methods for "LOXL2-dependent deacetylation of aldolase A induces metabolic reprogramming and tumor progression"

**Cells, transfections and mutagenesis**

Cells were maintained in either RPMI 1640 or DMEM medium supplemented with 10% fetal bovine serum and 100 mg/mL penicillin/streptomycin under standard conditions. All cell lines were authenticated and tested routinely for *Mycoplasma* contamination. All plasmids and specific siRNAs against human LOXL2 gene (siLOXL2#1: 5’-GCAACCGGCTCCTGAGTAT-3’; siLOXL2#2: 5’-CCAGATAGAGAACCTGAAT-3’; GenePharma) were transiently transfected into the indicated cells by using Lipofectamine 3000 (Life Technologies) according to the manufacturer’s instructions. KYSE510 cells infected with LOXL2 lentiviral shRNA (5’-TTCAACGGTGCTATAACCA-3’; Hanbio) and stable LOXL2-silenced KYSE510 cells with lentivirus expressing scrambled vector, Flag-tagged LOXL2 and L2Δ13 were obtained as described in our previous study(1).

Acetyl-mimetic and non-acetylatable mutants of aldolase A at lysine 13 were generated from pcDNA3.1-ALDOA-Flag by a Fast Mutagenesis System Kit (TransGen Biotech). The cDNAs encoding mutant LOXL2 constructs, including ND (1-544 aa), LOXL2-CD (545-774 aa) and L2Δ13-CD (545-729 aa), were amplified and cloned into the *Xho*I and *Hind*III sites in pcDNA3.1-HA (FM111-02, TransGen Biotech). Plasmid sequences were validated using Sanger sequencing. Details of primers are listed in Supplementary Table S5.

**Generation of L2Δ13-overexpressing mice**

For *L2Δ13*-overexpressing C57BL/6 transgenic mice, Flag-tagged human *L2Δ13* cDNA was inserted downstream to a *LoxP*-flanked *PGK-neomycin-polyA* cassette by homologous recombination into the *ROSA26* locus (Shanghai Biomodel Organism Co., Ltd). Conditional overexpressing mice were crossed with the CMV-Cre strain to obtain constitutive *L2Δ13* allele, and germ-line *L2Δ13*-overexpressing homozygous mice (Δ13/Δ13) were generated by intercrossing heterozygous mice (Δ13/-). *L2Δ13*-overexpressing ES cells and genotypes of mice were identified by PCR-based screening using PCR primers.

**Mouse models**

In diet-induced obesity mouse models, 16-week-old Δ13/Δ13 mice and littermate wild-type mice were fed either a high-fat diet (HFD, 60% energy in kcal from fat; D12492, Research DIETS) or normal chow diet (NCD, 10% energy in kcal from fat; D12450J, Research DIETS) for 8 weeks (n = 8-10 per group). Body weight was measured every week. Food intake and water intake per mouse was calculated weekly from the consumption per cage.

**Hematoxylin & eosin (HE) and immunohistochemistry**

Mouse tissue was fixed in 4% paraformaldehyde fixative prior to being embedded in paraffin. Human tissue microarrays were constructed as described previously(2). Fixed tissues were sectioned at 4 μm, and processed for H&E staining, and immunohistochemistry was performed using indicated antibodies following standard procedures.

**Oil Red O staining**

Pathological frozen sections of mouse livers were fixed in 10% formaldehyde, washed and dehydrated in 60% isopropanol. Oil Red O working solution (G1260, Solarbio Life Sciences) is prepared as follows: a saturated Oil Red O stock solution was freshly diluted to 60% with distilled water, mixed, equilibrated and filtered with a 0.22 μm-filter. Sections were then stained with the prepared Oil-Red-O stain for 15 min. After dehydration using isopropanol, samples were washed again and counterstained with hematoxylin solution. Evaluation of Oil Red O staining of each mouse was determined as positive or negative for further analysis.

**Periodic acid Schiff (PAS) staining**

Frozen livers of mice were cut into serial sections, fixed with Carnoy's Fluid, and rehydrated to deionized water for PAS staining (Leagene Biotechnology). The adjacent section of the same sample was treated with 1% amylase for 15 min at 37℃ and served as a negative control for glycogen staining. Frozen sections were incubated in 0.5% periodic acid, rinsed in water, and placed in Schiff’s reagent away from light. Samples were subsequently washed and counterstained with hematoxylin solution and dehydrated by consecutive washes in graded ethanol, followed by vitrification by dimethylbenzene. The level of PAS staining of individual tissue specimens was quantified according to its intensity, and the scores, which ranged from 0 to 4, were subjected to statistical analysis.

**Sirius red staining**

To detect collagen deposition, paraformaldehyde-fixed liver tissue was paraffin-embedded and sections were stained with a Sirius Red Stain Kit (DC0041, Leagene Biotechnology) according to the manufacturer’s recommendations. In brief, 4 μm sections were incubated with sirius red stain for 1 h and lightly washed in running water. After the counterstaining with Mayer’s hematoxylin for 10 min, the slides were washed with running water for 10 min. Sirius red-positive areas of each liver specimen were measured in at least three fields on each slide, and the staining was quantified using Image J (NIH) software.

**Blood chemistry analysis**

Liver dysfunction and kidney dysfunction were defined following standard liver and kidney biochemistry tests. Blood glucose, serum cholesterol (CHOL), low density lipoprotein cholesterol (LDL), triglyceride (TG), alanine aminotransferase (ALT) levels and kidney function indexes of mice (including blood urea nitrogen, creatinine and uric acid), were determined on an automated clinical chemistry analyzer (AU5831 series, Beckman Coulter) as recommended by the manufacturer (3).

**RNA-sequencing**

Total hepatic RNAs of wild-type and L2Δ13-overexpressing mice were isolated using Trizol reagent (15596018, Life Technology). Genome-wide RNA-sequencing with 6 biological replicates per group was performed using the BGISEQ-500 sequencing platform. Sequencing reads after filtering by SOAPnuke software were mapped to reference using HISAT (JHU) and Bowtie2 (http://bowtie-bio.sourceforge.net/bowtie2/index.shtml) tools, followed by strict quality control from different aspects. Gene expression was quantified with RSEM software. Differentially-expressed genes between groups were analyzed by screening using the DEGseq method according to a fold change ≥ 2. Pathway enrichment analysis of differentially-expressed genes was based on the KEGG database (http://www.genome.jp/kegg). The biological pathways with *p*-values of < 0.01 using were considered as significantly enriched functional annotations.

**Xenograft assays**

Xenograft assays in nude mice were performed as described previously(4). KYSE510 cells infected with scrambled shRNA (shNC) or shLOXL2 lentivirus (shLOXL2), or stable LOXL2-silenced KYSE510 cells expressing empty vector (vector), LOXL2-Flag (LOXL2) or L2Δ13-Flag (1 × 10^6^ cells in 100 μL PBS) were injected subcutaneously into the left flanks of BALB/c nude mice (Vital River Laboratories) at the age of 5 weeks ( n = 7 per group). General mouse behaviors were monitored and the tumor volume was measured every 3 days. The tumor size was calculated by the formula volume = 1/2 length × width^2^. Mice were euthanized after 30 days and tumor tissues were excised for growth analysis.

**Western blotting and co-immunoprecipitation**

For western blotting experiments, total protein lysates from cell lines and mouse livers were prepared in RIPA buffer, separated on 10% SDS-PAGE gels or the Stain-Free FastCast gels (1610183, Bio-Rad), transferred to PVDF membranes, and probed with the indicated primary antibodies against corresponding proteins and secondary HRP-conjugated antibodies. Immunoreactive bands were detected with a chemiluminescence detection system. Stain-Free FastCast gels, which served as protein loading controls, were activated and visualized using a Bio-Rad Stain-Free enabled imager (Bio-Rad)(5).

Co-immunoprecipitation (co-IP) assays were performed in HEK293T transfectants and stable KYSE510 cell lines. Briefly, 1 mg of whole cell lysate was immunoprecipitated overnight at 4℃ using Protein A/G PLUS-Agarose (sc-2003, Santa Cruz Biotechnology) pre-incubated with 1 μg anti-Flag antibody (F3165, Sigma-Aldrich), anti-HA antibody (sc-7392, Santa Cruz Biotechnology) or pan anti-acetyl lysine antibody (PTM-102, PTM Biolabs Inc). After three washes with PBS or TBS, the agarose precipitates were boiled in SDS-sample loading buffer prior to loading onto a gel for separation and PVDF membrane transfer for western blotting as described above.

**Pull-down assays**

KYSE510 cells or LOXL2-Flag/L2Δ13-Flag transfected HEK293T cells were harvested in bicine buffer (Thermo scientific). GST-Aldolase A fusion proteins were immobilized on glutathione resins, and then incubated with the cell lysates or purified recombinant Flag-tagged LOXL2/L2Δ13 at 4 °C for 2 h. Following precipitation, the cell pellets were washed 3 times with the lysis buffer (pH 7.3) containing 4.3 mM Na_2_HPO_4_, 1.47 mM KH_2_PO_4_, 137 mM NaCl and 0.1% Triton X-100, and ultimately analyzed by Western blotting with indicated antibodies.

***In Situ* proximity ligation assay (PLA) and confocal fluorescence microscopy**

Duolink *in situ* PLA (DUO92014, Sigma-Aldrich) was performed in esophageal cancer cells. The paired-primary antibodies used in the present study were either rabbit anti-LOXL2, L2Δ13 or GAPDH antibody with mouse anti-aldolase A antibody. As a negative control, PLA was performed using only anti-LOXL2, anti-L2Δ13 or anti-aldolase A antibody, respectively. Briefly, cells were fixed with 4% paraformaldehyde for 10 min, washed three times with PBS, and permeabilized in 0.1% Triton X-100 for 10 min, followed by incubation with the indicated antibody pairs overnight at 4 °C. PLA was performed according to the manufacturer’s recommendations.

To determine the co-localization of LOXL2 or L2Δ13 with aldolase A in esophageal cancer cells, immunofluorescence assays were performed in KYSE510 cells, which express these proteins endogenously, as previously described (2). To examine the effects of different aldolase A mutants on its mobilization, KYSE510 cells transfected with empty vector, Flag-tagged wild-type aldolase A, its Flag-tagged acetyl-mimetic or non-acetylatable mutant were stained with Flag antibody and Actin-stain™ 555 phalloidin, followed by counterstaining with DAPI for cell nuclei.

Confocal imaging was conducted on a laser-scanning confocal microscope (LSM 800, Carl Zeiss) with a 63 ×, 1.40 NA, Plan-apochromatic oil objective.

**Proliferation assays**

For colony formation assays, 1000 cells were seeded onto 6-well plates after 12 h of serum starvation, and were generally cultured for 2 weeks. Cells were fixed with 4% paraformaldehyde and stained with 0.5% crystal violet. The colony number of each group was calculated with Image J (NIH).

To monitor cell proliferation by directly measuring DNA synthesis, the Click‑iT^®^ Plus EdU Assay (C10637, Molecular probes) was conducted according to the manufacturer’s recommendations. Specifically, 5 × 10^4^ cells were inoculated on coverslips in a 24-well plate and cultured for 8 h. Cells were incubated with prepared 10 μM EdU solution for 2 h, fixed with 4% paraformaldehyde and permeabilized with 0.5% Triton^®^ X-100. Click-iT^®^ Plus reaction cocktail was applied to detect EdU of cells, while Hoechst^®^ 33342 solution (5 μg/mL) was used to for nuclear staining of cells. The EdU-labeled cells were ultimately observed and imaged on an Axio Observer A1 fluorescence microscope (Carl Zeiss) and analyzed quantifiably by Image J (NIH) software.

**ATP and glucose uptake assays**

Cells were seeded into 6-well plates and were transfected or infected with the indicated constructs. After 48 h, cells were harvested and 1×10^4^ cells were subsequently plated into a 96-well plate and cultured for 8 h. Cells were collected, extracted and incubated using the CellTiter-Glo^®^ Luminescent Cell Viability Assay (G7570, Promega) for ATP level analysis. Moreover, the Glucose Uptake-Glo™ Assay (J1342, Promega) was performed to measure glucose uptake in cells based on the detection of 2-deoxyglucose-6-phosphate (2DG6P). Luminescence was measured and recorded on a GloMax^®^ luminometer (Promega). The group of wells containing medium without cells served as negative controls.

**Lactate production assays**

For lactate production assays, cells were transfected or infected as for the ATP quantitation. After the determination of cell number, 5×10^4^ cells were plated into a 24-well plate and cultured for 12 h. To measure the secretion of lactate in the cell supernatants, the medium was collected and deproteinized with a 10 kDa MWCO spin filter to remove lactate dehydrogenase. After adding 200 μL Lactate Assay Buffer of the Lactate Assay Kit (MAK064, Sigma-Aldrich) to each sample, cell lysate was centrifuged at 13,000 × g for 10 min to remove insoluble material and then was deproteinized as described for the medium. Subsequently, 2.5 μL of prepared medium of cells and 10 μL of prepared cell lysis were brought to a final volume of 50 μL/well with Lactate Assay Buffer. To measure lactate production, each of the reaction content per well was incubated with 50 μL of Master Reaction Mix for 30 min at room temperature, protected from light. The reaction mixture was measured at 570 nm based on colorimetric assays.

**Aldolase enzyme activity assays**

For aldolase enzymatic activity of whole cell lysate, 1×10^6^ cells were homogenized in 100 μL of ice cold Aldolase Assay Buffer for 10 min and centrifuged at 10,000 × g at 4℃ for 5 min. The supernatant was analyzed by an Aldolase Activity Colorimetric Assay Kit (MAK223, Sigma-Aldrich). Samples were added into duplicate wells and incubated with the appropriate Reaction Mix at 37℃ for 1 to 60 min. Absorbance was measured at 450 nm in a microplate reader and was recorded in 1 min intervals.

For aldolase activity in the diffusible fraction, 8% of total supernatant (40 μL) and 10% of total cell lysate (20 μL) cell lysate were used for western blotting analysis. Another 40 μL supernatant was directly prepared for examining aldolase activity in the supernatant according to Boyer's modification of the hydrazine assay in which 3-phosphoglyceraldehyde reacts with hydrazine to generate a hydrazine(6). The cell supernatant was mixed with 6 μL of 0.01M iodoacetate, 3 μL of 0.01 M EDTA, 200 μL of 0.0035 M hydrazine, and an appropriate volume of lysis buffer to make a final volume of 300 μL. The Sample Blank readings were taken at 240 nm and were subtracted from the sample readings. Then, 10 μL of FBP (0.12 M) was added to each well and absorbance was detected at 240 nm every minute for 30 min.

**SILAC labeling, protein extraction, digestion and peptide enrichment**

KYSE510-shNC and stable LOXL2-depleted cells (KYSE510-shLOXL2) were grown for seven passages in RPMI 1640 medium for SILAC (88365, Thermo Fisher Scientific) supplemented with “heavy” (CLM-2247-H and CLM-2265-H, CIL) and “light” (L8662 and A8094, Sigma-Aldrich) arginine and lysine, and 10% fetal bovine serum, respectively. The cells were harvested, washed with cold PBS and further lysed by cell lysis buffer (1% protease inhibitor cocktail, 5 mM DTT, 2 mM EDTA, 3 μM TSA, 50 mM nicotinamide, pH 8.0). Protein extraction, digestion and fractionation were carried out according to our previous methods(7). Briefly, equal amounts of proteins from “heavy” labeled KYSE510-shNC cells and “light” labeled KYSE510-shLOXL2 cells were mixed, and then the protein mixture was reduced with 5 mM DTT at 56℃ for 30 min, and alkylated with 15 mM iodoacetamide for 45 min at room temperature in the dark, and further digested with trypsin. The obtained peptides were fractionated on high pH reversed phase HPLC into 10 fractions, followed by affinity enrichment of acetylated peptides using agarose-conjugated anti-acetyl-lysine antibody.

**Nano-LC-MS/MS**

Enriched acetylated peptides were analyzed by nano-HPLC-MS/MS as described previously(7,8). Specifically, each sample of peptides was separated on a 100 min gradient reversed-phase analytical column (Acclaim PepMap RSLC, Thermo Fisher Scientific). The resulting peptides were subjected to a nanospray ionization (NSI) source followed by tandem mass spectrometry (MS/MS) in a Q Exactive Plus Orbitrap mass spectrometer (Thermo Fisher Scientific). Intact peptides were detected in an Orbitrap analyzer (Thermo Fisher Scientific) at a resolution of 70,000. Peptides were selected for MS/MS using 30 as a normalized collision-energy (NCE) setting; ion fragments were detected in the Orbitrap at a resolution of 17,500. A data-dependent procedure that switched between one MS scan and 15 MS/MS scans (m/z 350-1800) was applied for the top 15 precursor ions.

**Database analysis**

The acquired MS/MS data was searched by MaxQuant (v.1.5.5.1) against the Uniport-Human protein sequence database (20,205 entries, http://www.uniprot.org) with an overall false discovery rate (FDR) for peptides of less than 1%(9). The MaxQuant database search was carried out with the following parameters: trypsin specificity, a maximum of 2 possible missed cleavages, 10 ppm mass tolerance for precursor ions, and 0.02 Da mass tolerance for MS/MS. The minimal peptide length was set to seven. Carbamidomethylation on Cys was specified as fixed modification. Acetylation on lysine, oxidation of methionine and acetylation on the peptide N-terminus were fixed as variable modifications. The quantification of acetyl peptides was normalized based on their corresponding protein levels, and differentially-acetylated proteins and sites were considered significant when the fold change was > 1.5.
