## Supplementary Tables S1-S6 for "LOXL2-dependent deacetylation of aldolase A induces metabolic reprogramming and tumor progression"

**Table S1. General characteristics and blood chemistry parameters of L2Δ13-overexpressing and age-matched wild-type mice.**

| **Index** | **4 weeks** | | |  | **8 weeks** | | |  | **16 weeks** | | |
| --- | --- | --- | --- | --- | --- | --- | --- | --- | --- | --- | --- |
|  | **WT** | **Δ13/Δ13** | ***P-*value** |  | **WT** | **Δ13/Δ13** | ***P-*value** |  | **WT** | **Δ13/Δ13** | ***P-*value** |
|  | **n = 19** | **n = 19** |  |  | **n = 20** | **n = 19** |  |  | **n = 20** | **n = 19** |  |
| **Body weight (g)** | 9.88 ± 2.01 | 9.33 ± 1.29 | 0.191 |  | 17.39 ± 1.95 | 16.43 ± 1.53 | 0.095 |  | 22.17 ± 3.07 | 19.86 ± 2.07 | 0.035* |
| **Body fat (%)** | 0.87 ± 0.23 | 0.64 ± 0.15 | 0.001*** |  | 2.55 ± 0.50 | 1.43 ± 0.27 | 0.001*** |  | 3.66 ± 1.00 | 2.70 ± 0.64 | 0.001*** |
| **BUN (mmol/L)** | 12.45 ± 2.33 | 11.7± 2.35 | 0.405 |  | 9.75 ± 1.94 | 11.87 ± 2.77 | 0.003** |  | 10.15 ± 1.55 | 10.35 ± 2.27 | 0.967 |
| **CHE (U/mL）** | / | / | / |  | 5.04 ± 0.66 | 6.82 ± 0.73 | < 0.001*** |  | 4.59 ± 0.89 | 5.58 ± 0.62 | 0.001*** |
| **CHOL (mmol/L)** | 1.55 ± 0.37 | 1.83 ± 0.35 | 0.026* |  | 1.63 ± 0.29 | 2.81 ± 0.40 | 0.001*** |  | 2.29 ± 0.39 | 2.99 ± 0.57 | 0.001*** |
| **Creatinine (umol/L)** | 24 ± 3.23 | 21.83 ± 2.17 | 0.051 |  | 40.50 ± 3.50 | 44.89 ± 1.76 | < 0.001 |  | 46.20 ± 10.13 | 44.26 ± 9.13 | 0.411 |
| **Glucose (mmol/L)** | 3.59 ± 1.11 | 2.85 ± 0.54 | 0.032* |  | 6.24 ± 1.99 | 4.61 ± 0.94 | 0.008** |  | 6.96 ± 0.96 | 5.11 ± 1.27 | 0.001*** |
| **LDL (mmol/L）** | 0.34 ± 0.09 | 0.48 ± 0.11 | 0.001*** |  | 0.45 ± 0.08 | 0.64 ± 0.09 | 0.001*** |  | 0.46 ± 0.08 | 0.69 ± 0.18 | 0.001*** |
| **TG (mmol/L)** | 0.88 ± 0.34 | 1.12 ± 0.36 | 0.043* |  | 1.38 ± 0.34 | 2.23 ± 0.63 | 0.001*** |  | 0.89 ± 0.24 | 1.13 ± 0.36 | 0.019* |
| **Uric acid (umol/L)** | 74.00 ± 28.17 | 58.74 ± 14.50 | 0.116 |  | 32.85 ± 7.57 | 33.00 ± 4.45 | 0.444 |  | 34.55 ± 7.15 | 33.16 ± 4.90 | 0.945 |
| BUN, blood urea nitrogen; CHE, cholinesterase; CHOL, cholesterol; LDL, low density lipoprotein cholesterol; TG, triglycerides. | | | | | | | | | | | |
| The *p*-value was calculated by unpaired Student's *t* test or nonparametric test. | | | | | |  |  |  |  |  |  |

**Table S2. Lysine acetylation of proteins regulated by LOXL2 and L2Δ13 in esophageal cancer cells.**

| **Gene name** | **Ratio shLOXL2/shNC** | **Protein ID** | **Position** |
| --- | --- | --- | --- |
| ACAD11 | 6.24 | Q709F0 | 704 |
| ACAD9 | 3.39 | Q9H845 | 294 |
| ACADM | 5.75 | P11310 | 279 |
| ACADM | 1.04 | P11310 | 271 |
| ACAT1 | 2.46 | P24752 | 174 |
| ACOT4 | NA | Q8N9L9 | 415 |
| ACOT8 | 9.38 | O14734 | 169 |
| ACOT9 | 2.78 | Q9Y305 | 407 |
| ACOX1 | 9.20 | Q15067 | 267 |
| ACOX1 | 8.51 | Q15067 | 500 |
| ACOX1 | 6.37 | Q15067 | 437 |
| ACOX1 | 3.38 | Q15067 | 512 |
| ACSL1 | 0.93 | P33121 | 207 |
| ACTA1;ACTC1;ACTG2;ACTA2 | 1.75 | P68133;P68032;P63267;P62736 | 70;70;69;70 |
| ACTB;ACTG1;ACTA1;ACTC1;ACTG2;ACTA2 | 1.30 | P60709;P63261;P68133;P68032;P63267;P62736 | 326;326;328;328;327;328 |
| ACTB;ACTG1;ACTA1;ACTC1;ACTG2;ACTA2 | 1.10 | P60709;P63261;P68133;P68032;P63267;P62736 | 61;61;63;63;62;63 |
| ADNP | NA | Q9H2P0 | 716 |
| AHNAK | NA | Q09666 | 134 |
| AHNAK | 5.99 | Q09666 | 730 |
| AHNAK | 2.13 | Q09666 | 609 |
| AHNAK2 | 1.07 | Q8IVF2 | 1932 |
| AHNAK2 | 1.07 | Q8IVF2 | 1936 |
| AIM1 | 14.98 | Q9Y4K1 | 308 |
| AIM1 | 14.98 | Q9Y4K1 | 313 |
| AKR1C2;AKR1C1 | 1.69 | P52895;Q04828 | 185;185 |
| AKR1C2;AKR1C1;AKR1C3;AKR1C4 | 0.74 | P52895;Q04828;P42330;P17516 | 270;270;270;270 |
| ALDH6A1 | 2.20 | Q02252 | 129 |
| ALDOA | 1.62 | P04075 | 13 |
| ALDOA | 1.38 | P04075 | 230 |
| ALDOA | 1.14 | P04075 | 42 |
| ANXA1 | 0.85 | P04083 | 312 |
| ANXA2 | 1.36 | P07355 | 49 |
| ANXA2 | 1.33 | P07355 | 266 |
| APEX1 | NA | P27695 | 24 |
| ARID1B | 5.48 | Q8NFD5 | 648 |
| ASCC3 | 1.58 | Q8N3C0 | 572 |
| ATP1A1 | 2.43 | P05023 | 9 |
| ATP2A2 | 1.07 | P16615 | 628 |
| ATP5A1 | NA | P25705 | 230 |
| ATP5A1 | 1.83 | P25705 | 531 |
| ATP5A1 | 1.44 | P25705 | 539 |
| ATP5F1 | 1.55 | P24539 | 162 |
| ATRX | 2.82 | P46100 | 967 |
| BAIAP3 | 1.48 | O94812 | 308 |
| BCAT2 | 2.37 | O15382 | 141 |
| BRD2 | NA | P25440 | 31 |
| C1QBP | 1.19 | Q07021 | 91 |
| CARD17;CARD16;CASP1 | 1.09 | Q5XLA6;Q5EG05;P29466 | 7;7;7 |
| CARS | 72.87 | P49589 | 412 |
| CARS | 72.87 | P49589 | 418 |
| CAT | 7.61 | P04040 | 480 |
| CAT | 7.53 | P04040 | 93 |
| CAT | 3.70 | P04040 | 243 |
| CAV1 | 2.26 | Q03135 | 5 |
| CBX3 | NA | Q13185 | 103 |
| CBX3 | 1.52 | Q13185 | 5 |
| CCDC136 | 34.71 | Q96JN2 | 549 |
| CCDC136 | 34.71 | Q96JN2 | 551 |
| CCNYL1;CCNY | 0.88 | Q8N7R7;Q8ND76 | 94;72 |
| CCT6A | 1.75 | P40227 | 5 |
| CCT6A | 1.13 | P40227 | 10 |
| CCT6A | 1.01 | P40227 | 377 |
| CHAMP1 | 3.32 | Q96JM3 | 617 |
| CIC | 6.87 | Q96RK0 | 689 |
| CNBP | NA | P62633 | 8 |
| CRAT | 7.43 | P43155 | 609 |
| CREBBP | 3.62 | Q92793 | 1239 |
| CREBBP | 2.96 | Q92793 | 1216 |
| CREBBP | 2.75 | Q92793 | 1014 |
| CREBBP | 2.32 | Q92793 | 1586 |
| CREBBP | 2.12 | Q92793 | 1583 |
| CREBBP | 2.11 | Q92793 | 1736 |
| CREBBP | 2.06 | Q92793 | 1595 |
| CREBBP | 2.06 | Q92793 | 1597 |
| CREBBP | 0.17 | Q92793 | 13 |
| CS | 1.73 | O75390 | 76 |
| CSNK1A1 | NA | P48729 | 8 |
| CTCF | 1.77 | P49711 | 18 |
| CTTN | NA | Q14247 | 171 |
| CTTN | 2.32 | Q14247 | 107 |
| CTTN | 1.82 | Q14247 | 205 |
| CTTN | 1.66 | Q14247 | 295 |
| CTTN | 1.53 | Q14247 | 110 |
| CTTN | 1.49 | Q14247 | 319 |
| CTTN | 1.45 | Q14247 | 181 |
| CTTN | 1.42 | Q14247 | 208 |
| CTTN | 1.37 | Q14247 | 245 |
| CTTN | 1.30 | Q14247 | 144 |
| CTTN | 1.02 | Q14247 | 198 |
| CTTN | 0.96 | Q14247 | 87 |
| CTTN | 0.92 | Q14247 | 346 |
| CTTN | 0.90 | Q14247 | 161 |
| CTTN | 0.80 | Q14247 | 272 |
| CTTN | 0.78 | Q14247 | 124 |
| CTTN | 0.75 | Q14247 | 235 |
| CYTH4 | 1.06 | Q9UIA0 | 108 |
| DBI | 1.63 | P07108 | 19 |
| DBI | 1.09 | P07108 | 55 |
| DBT | 1.95 | P11182 | 440 |
| DCPS | 2.43 | Q96C86 | 10 |
| DDX18 | 1.68 | Q9NVP1 | 458 |
| DDX24 | 3.03 | Q9GZR7 | 17 |
| DDX39B | 2.00 | Q13838 | 36 |
| DDX42 | 0.86 | Q86XP3 | 145 |
| DDX46 | 0.58 | Q7L014 | 776 |
| DDX47 | 1.48 | Q9H0S4 | 161 |
| DECR2 | 2.30 | Q9NUI1 | 29 |
| DHCR7 | 4.99 | Q9UBM7 | 4 |
| DKC1 | 3.45 | O60832 | 11 |
| DLD | NA | P09622 | 166 |
| DLD | 2.62 | P09622 | 159 |
| DLD | 2.22 | P09622 | 143 |
| DLD | 1.66 | P09622 | 122 |
| DLD | 1.66 | P09622 | 430 |
| DNPEP | 11.78 | Q9ULA0 | 51 |
| ECH1 | 5.42 | Q13011 | 250 |
| ECH1 | 4.63 | Q13011 | 112 |
| ECH1 | 2.74 | Q13011 | 196 |
| ECH1 | 2.60 | Q13011 | 95 |
| ECHS1 | 1.53 | P30084 | 118 |
| ECI1 | 1.53 | P42126 | 283 |
| ECI2 | 9.12 | O75521 | 62 |
| ECI2 | 7.90 | O75521 | 92 |
| ECI2 | 7.18 | O75521 | 51 |
| ECI2 | 6.74 | O75521 | 305 |
| ECI2 | 4.45 | O75521 | 161 |
| EEF2 | 1.53 | P13639 | 239 |
| EIF3L | 0.54 | Q9Y262 | 549 |
| EIF4B | 0.64 | P23588 | 13 |
| EIF5A;EIF5A2 | 1.73 | P63241;Q9GZV4 | 47;47 |
| ENO1 | 1.09 | P06733 | 193 |
| ENO1 | 0.99 | P06733 | 5 |
| ENO1 | 0.74 | P06733 | 343 |
| EP300 | 10.08 | Q09472 | 14 |
| EP300 | 3.72 | Q09472 | 1203 |
| EP300 | 2.59 | Q09472 | 1546 |
| EP300 | 2.16 | Q09472 | 1180 |
| EP300 | 1.88 | Q09472 | 1542 |
| EP300 | 1.48 | Q09472 | 1558 |
| EP300 | 1.48 | Q09472 | 1560 |
| EP300;CREBBP | 3.04 | Q09472;Q92793 | 1499;1535 |
| EP300;CREBBP | 2.67 | Q09472;Q92793 | 1769;1806 |
| EP300;CREBBP | 2.67 | Q09472;Q92793 | 1772;1809 |
| EP300;CREBBP | 2.20 | Q09472;Q92793 | 1674;1711 |
| EPHA5 | 0.91 | P54756 | 1018 |
| EXOSC10 | 1.92 | Q01780 | 102 |
| FAAH2 | NA | Q6GMR7 | 429 |
| FASN | 1.26 | P49327 | 70 |
| FASN | 0.93 | P49327 | 1771 |
| FASN | 0.92 | P49327 | 1072 |
| FASN | 0.87 | P49327 | 2449 |
| FASN | 0.86 | P49327 | 1752 |
| FASN | 0.77 | P49327 | 1704 |
| FAU | NA | P62861 | 51 |
| FH | 2.74 | P07954 | 66 |
| FLNA | 1.04 | P21333 | 700 |
| FSCN2 | 2.49 | O14926 | 41 |
| G6PD | 4.37 | P11413 | 497 |
| GAPDH | 1.23 | P04406 | 194 |
| GFM2 | 0.78 | Q969S9 | 83 |
| GLUD1 | 1.92 | P00367 | 84 |
| GLUD1 | 1.66 | P00367 | 503 |
| GMNN | 2.08 | O75496 | 27 |
| GOT2 | 2.54 | P00505 | 73 |
| GOT2 | 1.80 | P00505 | 185 |
| GOT2 | 1.75 | P00505 | 122 |
| GOT2 | 1.70 | P00505 | 309 |
| GOT2 | 1.66 | P00505 | 90 |
| GOT2 | 1.46 | P00505 | 159 |
| GPD2 | 4.27 | P43304 | 436 |
| GSTK1 | 11.87 | Q9Y2Q3 | 54 |
| GSTK1 | 10.26 | Q9Y2Q3 | 71 |
| GTF2I | 7.40 | P78347 | 353 |
| H1FX | 2.83 | Q92522 | 19 |
| H2AFV;H2AFZ | NA | Q71UI9;P0C0S5 | 116;116 |
| H2AFV;H2AFZ | 2.07 | Q71UI9;P0C0S5 | 5;5 |
| H2AFV;H2AFZ | 2.07 | Q71UI9;P0C0S5 | 8;8 |
| H2AFY | NA | O75367 | 116 |
| HADHA | 2.45 | P40939 | 597 |
| HADHA | 1.66 | P40939 | 644 |
| HADHA | 1.54 | P40939 | 406 |
| HADHA | 0.83 | P40939 | 505 |
| HADHB | 2.61 | P55084 | 72 |
| HADHB | 2.07 | P55084 | 111 |
| HAT1 | 0.67 | O14929 | 9 |
| HDAC2 | 0.82 | Q92769 | 452 |
| HDGF | 1.63 | P51858 | 39 |
| HDHD3 | 1.85 | Q9BSH5 | 15 |
| HIST1H1B;HIST1H1C;HIST1H1E;HIST1H1D;HIST1H1A | NA | P16401;P16403;P10412;P16402;Q02539 | 93;90;90;91;93 |
| HIST1H1B;HIST1H1C;HIST1H1E;HIST1H1D;HIST1H1A | 1.76 | P16401;P16403;P10412;P16402;Q02539 | 100;97;97;98;100 |
| HIST1H2AJ;HIST1H2AH;H2AFJ;HIST1H2AD;HIST1H2AG;HIST1H2AC;HIST3H2A;HIST1H2AB;H2AFX;HIST2H2AC;HIST2H2AA3 | NA | Q99878;Q96KK5;Q9BTM1;P20671;P0C0S8;Q93077;Q7L7L0;P04908;P16104;Q16777;Q6FI13 | 119;119;119;119;119;119;119;119;119;119;119 |
| HIST1H2BB | 1.33 | P33778 | 17 |
| HIST1H2BB | 1.33 | P33778 | 21 |
| HIST1H2BC;H2BFS;HIST1H2BK | 30.87 | P62807;P57053;O60814 | 6;6;6 |
| HIST1H2BC;H2BFS;HIST1H2BK | 0.85 | P62807;P57053;O60814 | 12;12;12 |
| HIST1H2BC;H2BFS;HIST1H2BK;HIST1H2BN;HIST1H2BD;HIST1H2BA;HIST1H2BO;HIST1H2BB;HIST2H2BF;HIST1H2BH;HIST1H2BM;HIST1H2BL | 3.15 | P62807;P57053;O60814;Q99877;P58876;Q96A08;P23527;P33778;Q5QNW6;Q93079;Q99879;Q99880 | 109;109;109;109;109;110;109;109;109;109;109;109 |
| HIST1H2BC;H2BFS;HIST1H2BK;HIST1H2BN;HIST1H2BD;HIST1H2BO;HIST1H2BH;HIST1H2BL | 2.54 | P62807;P57053;O60814;Q99877;P58876;P23527;Q93079;Q99880 | 17;17;17;17;17;17;17;17 |
| HIST1H2BC;H2BFS;HIST1H2BK;HIST1H2BN;HIST1H2BD;HIST1H2BO;HIST1H2BH;HIST1H2BL | 2.05 | P62807;P57053;O60814;Q99877;P58876;P23527;Q93079;Q99880 | 21;21;21;21;21;21;21;21 |
| HIST1H2BC;H2BFS;HIST1H2BK;HIST1H2BN;HIST1H2BD;HIST1H2BO;HIST1H2BH;HIST1H2BL | 1.66 | P62807;P57053;O60814;Q99877;P58876;P23527;Q93079;Q99880 | 24;24;24;24;24;24;24;24 |
| HIST1H2BL | 5.60 | Q99880 | 6 |
| HIST1H2BM | 7.44 | Q99879 | 6 |
| HIST1H2BM | 2.38 | Q99879 | 17 |
| HIST1H2BM | 2.38 | Q99879 | 21 |
| HIST1H2BO;HIST2H2BF;HIST1H2BH | 8.99 | P23527;Q5QNW6;Q93079 | 12;12;12 |
| HIST1H2BO;HIST2H2BF;HIST1H2BH | 2.58 | P23527;Q5QNW6;Q93079 | 6;6;6 |
| HIST1H3A;H3F3A;HIST3H3;HIST2H3A | NA | P68431;P84243;Q16695;Q71DI3 | 80;80;80;80 |
| HIST1H3A;H3F3C;H3F3A;HIST2H3A | 2.44 | P68431;Q6NXT2;P84243;Q71DI3 | 24;24;24;24 |
| HIST1H3A;H3F3C;H3F3A;HIST2H3A | 2.18 | P68431;Q6NXT2;P84243;Q71DI3 | 19;19;19;19 |
| HIST1H3A;H3F3C;H3F3A;HIST3H3;HIST2H3A | 3.08 | P68431;Q6NXT2;P84243;Q16695;Q71DI3 | 57;56;57;57;57 |
| HIST1H3A;H3F3C;H3F3A;HIST3H3;HIST2H3A | 2.81 | P68431;Q6NXT2;P84243;Q16695;Q71DI3 | 10;10;10;10;10 |
| HIST1H3A;H3F3C;H3F3A;HIST3H3;HIST2H3A | 2.81 | P68431;Q6NXT2;P84243;Q16695;Q71DI3 | 15;15;15;15;15 |
| HIST1H3A;HIST3H3;HIST2H3A | 5.19 | P68431;Q16695;Q71DI3 | 28;28;28 |
| HIST1H4A | NA | P62805 | 92 |
| HIST1H4A | 2.73 | P62805 | 17 |
| HIST1H4A | 2.71 | P62805 | 13 |
| HIST1H4A | 2.68 | P62805 | 9 |
| HIST2H2BF | 2.23 | Q5QNW6 | 17 |
| HIST2H2BF | 2.23 | Q5QNW6 | 21 |
| HIST3H3 | 1.74 | Q16695 | 19 |
| HIST3H3 | 1.74 | Q16695 | 24 |
| HMGA1 | 1.60 | P17096 | 15 |
| HMGA1 | 1.41 | P17096 | 7 |
| HMGCL | 7.38 | P35914 | 93 |
| HMGCL | 5.99 | P35914 | 48 |
| HMGN1 | NA | P05114 | 14 |
| HMGN2 | 2.50 | P05204 | 82 |
| HNRNPA1 | 2.23 | P09651 | 3 |
| HNRNPA1 | 2.05 | P09651 | 350 |
| HNRNPA1;HNRNPA1L2 | 1.43 | P09651;Q32P51 | 105;105 |
| HNRNPD | 2.62 | Q14103 | 218 |
| HNRNPD | 2.26 | Q14103 | 231 |
| HNRNPDL | 2.51 | O14979 | 269 |
| HNRNPK | 1.89 | P61978 | 405 |
| HNRNPL | 4.14 | P14866 | 62 |
| HNRNPM | 2.19 | P52272 | 698 |
| HNRNPU | NA | Q00839 | 565 |
| HNRNPU | 4.64 | Q00839 | 186 |
| HNRNPU | 3.43 | Q00839 | 9 |
| HNRNPU | 1.65 | Q00839 | 352 |
| HSD17B4 | 12.99 | P51659 | 403 |
| HSD17B4 | 12.96 | P51659 | 415 |
| HSD17B4 | 11.33 | P51659 | 565 |
| HSD17B4 | 9.28 | P51659 | 84 |
| HSD17B4 | 8.63 | P51659 | 663 |
| HSD17B4 | 6.85 | P51659 | 46 |
| HSD17B4 | 6.17 | P51659 | 621 |
| HSD17B4 | 4.95 | P51659 | 57 |
| HSD17B4 | 1.26 | P51659 | 424 |
| HSDL2 | 11.69 | Q6YN16 | 116 |
| HSDL2 | 6.16 | Q6YN16 | 42 |
| HSDL2 | 5.18 | Q6YN16 | 49 |
| HSP90AA1;HSP90AA2P;HSP90AB1;HSP90AB2P | 1.21 | P07900;Q14568;P08238;Q58FF8 | 283;282;275;197 |
| HSP90AB1 | 1.41 | P08238 | 624 |
| HSPA1B;HSPA1A | 1.64 | P0DMV9;P0DMV8 | 56;56 |
| HSPA1B;HSPA1A | 1.08 | P0DMV9;P0DMV8 | 3;3 |
| HSPA1B;HSPA1A;HSPA6 | NA | P0DMV9;P0DMV8;P17066 | 500;500;502 |
| HSPA1B;HSPA1A;HSPA6 | 1.05 | P0DMV9;P0DMV8;P17066 | 507;507;509 |
| HSPA4 | 0.76 | P34932 | 53 |
| HSPA5 | NA | P11021 | 213 |
| HSPA5 | 4.61 | P11021 | 81 |
| HSPA5 | 2.22 | P11021 | 326 |
| HSPA8;HSPA2 | NA | P11142;P54652 | 187;188 |
| HSPA8;HSPA2 | 1.05 | P11142;P54652 | 507;510 |
| HSPA9 | 2.16 | P38646 | 600 |
| HSPA9 | 1.56 | P38646 | 394 |
| HSPA9 | 1.23 | P38646 | 377 |
| HSPD1 | 1.68 | P10809 | 58 |
| HSPD1 | 1.59 | P10809 | 481 |
| HSPD1 | 1.55 | P10809 | 269 |
| HSPD1 | 1.24 | P10809 | 205 |
| HSPD1 | 1.21 | P10809 | 156 |
| HSPE1 | NA | P61604 | 8 |
| HSPE1 | 1.58 | P61604 | 80 |
| HSPE1 | 1.10 | P61604 | 40 |
| HSPE1 | 1.10 | P61604 | 86 |
| HYPK | 1.08 | Q9NX55 | 35 |
| IARS2 | 1.97 | Q9NSE4 | 233 |
| IARS2 | 1.79 | Q9NSE4 | 803 |
| IDH1 | 2.24 | O75874 | 224 |
| IDH2 | 5.87 | P48735 | 80 |
| IFI16 | 3.52 | Q16666 | 128 |
| IFIT3 | NA | O14879 | 6 |
| ILF3 | NA | Q12906 | 600 |
| ILF3 | 1.87 | Q12906 | 460 |
| ING4 | 6.33 | Q9UNL4 | 146 |
| ING4 | 6.33 | Q9UNL4 | 148 |
| ING4 | 3.34 | Q9UNL4 | 127 |
| ING4 | 3.34 | Q9UNL4 | 129 |
| INTS3 | 5.29 | Q68E01 | 986 |
| INTS3 | 2.87 | Q68E01 | 915 |
| INTS4 | 3.93 | Q96HW7 | 26 |
| ISOC2 | 2.09 | Q96AB3 | 26 |
| JADE2 | 4.56 | Q9NQC1 | 38 |
| JADE2 | 2.84 | Q9NQC1 | 32 |
| JADE3 | NA | Q92613 | 30 |
| JADE3 | NA | Q92613 | 32 |
| KAT5 | 6.30 | Q92993 | 104 |
| KAT6A | 2.06 | Q92794 | 815 |
| KAT7 | 19.13 | O95251 | 171 |
| KDM5B | 4.44 | Q9UGL1 | 832 |
| KDM6A | 4.26 | O15550 | 799 |
| KIAA1598 | NA | A0MZ66 | 230 |
| KIFC1 | NA | Q9BW19 | 233 |
| KMT2B | 12.27 | Q9UMN6 | 926 |
| KMT2C | NA | Q8NEZ4 | 758 |
| KMT2C | 11.08 | Q8NEZ4 | 2809 |
| KMT2C | 11.08 | Q8NEZ4 | 2814 |
| KMT2C | 3.33 | Q8NEZ4 | 1508 |
| KMT2D | 4.66 | O14686 | 3079 |
| KMT2D | 4.37 | O14686 | 3573 |
| KMT2D | 3.69 | O14686 | 3119 |
| KRT1 | 20.16 | P04264;CON__P04264 | 211;211 |
| KRT1 | 13.61 | P04264;CON__P04264 | 395;395 |
| KRT1 | 7.36 | P04264;CON__P04264 | 197;197 |
| KRT1 | 6.20 | P04264 | 364 |
| KRT10 | 33.77 | CON__P13645;P13645 | 207;207 |
| KRT10 | 10.48 | CON__P13645;P13645 | 266;266 |
| KRT17 | NA | CON__Q04695;Q04695;CON__Q9QWL7 | 419;419;420 |
| KRT19;KRT10;KRT15;KRT17 | 23.10 | CON__P08727;P08727;CON__P13645;P13645;CON__P19012;P19012;CON__Q04695;Q04695;CON__Q9QWL7 | 97;97;163;163;122;122;101;101;101 |
| KRT2 | 81.38 | CON__P35908;P35908 | 308;302 |
| KRT6C;KRT6B;KRT6A;KRT2 | NA | CON__P48668;CON__P04259;CON__P02538;P48668;P04259;P02538;CON__P35908;P35908 | 338;338;338;338;338;338;359;353 |
| KRT6C;KRT6B;KRT6A;KRT8;KRT5;KRT76;KRT2;KRT77;KRT3;KRT1 | 23.29 | CON__P48668;CON__P04259;CON__P02538;P48668;P04259;P02538;CON__P05787;P05787;CON__H-INV:HIT000016045;CON__P13647;P13647;CON__Q01546;Q01546;CON__Q3TTY5;CON__P35908;P35908;CON__Q7Z794;Q7Z794;CON__P50446;CON__P12035;P12035;CON__Q5XQN5;CON__Q922U2;CON__Q8VED5;P04264;CON__P04264 | 259;259;259;259;259;259;185;185;18;264;264;279;279;295;280;274;260;260;248;296;296;265;258;236;276;276 |
| KRT6C;KRT6B;KRT6A;KRT8;KRT84;KRT5;KRT76;KRT2;KRT3;KRT7 | 2.64 | CON__P48668;CON__P04259;CON__P02538;P48668;P04259;P02538;CON__P05787;P05787;CON__Q9H552;CON__Q9NSB2;CON__Q6ISB0;Q9NSB2;CON__P08729;CON__Q9DCV7;CON__P13647;P13647;CON__Q01546;Q01546;CON__P07744;CON__Q3TTY5;CON__P35908;P35908;CON__P50446;CON__P12035;P12035;CON__Q3KNV1;P08729;CON__Q5XQN5;CON__Q922U2;CON__Q8VED5 | 173;173;173;173;173;173;101;101;117;175;175;175;101;95;178;178;193;193;156;209;194;188;162;208;208;101;101;179;172;149 |
| KRT7 | 2.00 | CON__P08729;CON__Q3KNV1;P08729 | 130;130;130 |
| KTN1 | 2.51 | Q86UP2 | 272 |
| LACTB2 | 7.30 | Q53H82 | 102 |
| LACTB2 | 2.67 | Q53H82 | 247 |
| LARS | 0.88 | Q9P2J5 | 970 |
| LASP1 | 1.49 | Q14847 | 96 |
| LDHA | 2.82 | P00338 | 14 |
| LMNA | NA | P02545 | 450 |
| LMNA | 2.45 | P02545 | 417 |
| LMNA | 1.36 | P02545 | 311 |
| LMNB1 | 1.42 | P20700 | 271 |
| LRPPRC | 1.23 | P42704 | 750 |
| MAGOHB;MAGOH | 2.84 | Q96A72;P61326 | 18;16 |
| MATR3 | 6.19 | P43243 | 3 |
| MBD2 | NA | Q9UBB5 | 382 |
| MCCC1 | 4.81 | Q96RQ3 | 295 |
| MCM3 | 1.70 | P25205 | 556 |
| MCM4 | 1.49 | P33991 | 858 |
| MDH2 | 2.19 | P40926 | 185 |
| MDH2 | 1.46 | P40926 | 296 |
| ME2 | 1.62 | P23368 | 272 |
| ME2 | 1.41 | P23368 | 240 |
| MEAF6 | NA | Q9HAF1 | 91 |
| MEAF6 | 1.22 | Q9HAF1 | 69 |
| MEAF6 | 1.22 | Q9HAF1 | 74 |
| MED1 | 5.89 | Q15648 | 1076 |
| MED1 | 3.03 | Q15648 | 1152 |
| MED1 | 1.37 | Q15648 | 1354 |
| MED27 | 1.71 | Q6P2C8 | 145 |
| MED6 | NA | O75586 | 236 |
| MED6 | NA | O75586 | 241 |
| MPHOSPH6 | NA | Q99547 | 127 |
| MRPL39 | 1.62 | Q9NYK5 | 72 |
| MRPL47 | 1.41 | Q9HD33 | 144 |
| MTHFD1L | 1.04 | Q6UB35 | 189 |
| MTHFD2 | 0.98 | P13995 | 50 |
| MYBBP1A | 1.91 | Q9BQG0 | 190 |
| MYC | 0.95 | P01106 | 148 |
| MYO1C | 4.93 | O00159 | 361 |
| NAA30 | 1.52 | Q147X3 | 104 |
| NAA30 | 1.19 | Q147X3 | 233 |
| NAA50 | 1.33 | Q9GZZ1 | 34 |
| NACA | 1.08 | Q13765;E9PAV3 | 142;2005 |
| NAE1 | 0.86 | Q13564 | 6 |
| NANS | 0.89 | Q9NR45 | 355 |
| NAT10 | NA | Q9H0A0 | 426 |
| NCOA2 | 6.24 | Q15596 | 31 |
| NDUFAB1 | 2.70 | O14561 | 151 |
| NHP2L1 | NA | P55769 | 21 |
| NME2P1;NME2 | 0.83 | O60361;P22392 | 109;124 |
| NOLC1 | 22.12 | Q14978 | 510 |
| NOLC1 | 2.62 | Q14978 | 76 |
| NOLC1 | 2.31 | Q14978 | 579 |
| NONO | 4.31 | Q15233 | 198 |
| NONO | 2.17 | Q15233 | 5 |
| NOP2 | 1.97 | P46087 | 91 |
| NOP56 | NA | O00567 | 578 |
| NOP56 | 2.89 | O00567 | 533 |
| NOP58 | NA | Q9Y2X3 | 441 |
| NPAT | 2.60 | Q14207 | 543 |
| NPM1 | 3.00 | P06748 | 150 |
| NPM1 | 1.86 | P06748 | 267 |
| NPM1 | 1.75 | P06748 | 141 |
| NPM1 | 1.39 | P06748 | 32 |
| NUDT12 | 9.20 | Q9BQG2 | 123 |
| NUDT21 | 2.30 | O43809 | 23 |
| NUP153 | 2.68 | P49790 | 15 |
| NUP153 | 2.29 | P49790 | 227 |
| NUP153 | 1.66 | P49790 | 705 |
| NUP85 | 1.02 | Q9BW27 | 92 |
| OGDH | NA | Q02218 | 401 |
| OXCT1 | 1.77 | P55809 | 421 |
| PAICS | 0.87 | P22234 | 11 |
| PAPSS2 | 8.97 | O95340 | 523 |
| PARN | 1.68 | O95453 | 566 |
| PAXIP1 | 5.12 | Q6ZW49 | 847 |
| PDCD11 | 2.49 | Q14690 | 1363 |
| PDHA1 | 2.20 | P08559 | 321 |
| PDHA1 | 1.76 | P08559 | 336 |
| PDHB | 1.38 | P11177 | 68 |
| PDS5A | 1.71 | Q29RF7 | 1211 |
| PEX2 | 7.54 | P28328 | 84 |
| PGAM1;PGAM4;PGAM2 | 1.15 | P18669;Q8N0Y7;P15259 | 106;106;106 |
| PGAM1;PGAM4;PGAM2 | 1.04 | P18669;Q8N0Y7;P15259 | 100;100;100 |
| PGK1 | 2.05 | P00558 | 30 |
| PGK1 | 0.97 | P00558 | 131 |
| PHF5A | 2.07 | Q7RTV0 | 3 |
| PKM | 1.32 | P14618 | 115 |
| PKM | 1.26 | P14618 | 367 |
| PNPT1 | 1.34 | Q8TCS8 | 246 |
| PNPT1 | 0.91 | Q8TCS8 | 306 |
| PPIA | 2.05 | P62937 | 82 |
| PPIA | 2.05 | P62937 | 125 |
| PPIA | 1.33 | P62937 | 76 |
| PPM1G | NA | O15355 | 519 |
| PPM1G | 1.02 | O15355 | 383 |
| PRCC | 1.27 | Q92733 | 250 |
| PRKDC | 3.58 | P78527 | 2908 |
| PRKDC | 1.06 | P78527 | 838 |
| PSMA3 | 5.70 | P25788 | 57 |
| PSMC6 | 0.85 | P62333 | 206 |
| PSPC1 | 4.11 | Q8WXF1 | 519 |
| PTBP1 | 3.23 | P26599 | 13 |
| PTMA | 0.91 | P06454 | 15 |
| PYGL | 0.10 | P06737 | 29 |
| RAN | 1.63 | P62826 | 134 |
| RAN | 0.95 | P62826 | 71 |
| RANBP2 | 2.34 | P49792 | 1851 |
| RBBP4 | 2.08 | Q09028 | 4 |
| RBBP7 | 1.66 | Q16576 | 119 |
| RBBP7 | 1.45 | Q16576 | 4 |
| RBM14 | 0.95 | Q96PK6 | 149 |
| RBM33 | 2.34 | Q96EV2 | 23 |
| RCC2 | 2.54 | Q9P258 | 77 |
| RPA1 | 0.73 | P27694 | 167 |
| RPL24 | 2.44 | P83731 | 93 |
| RPL24 | 2.06 | P83731 | 27 |
| RPL27 | 0.86 | P61353 | 27 |
| RPL4 | 1.51 | P36578 | 364 |
| RPN1 | 1.19 | P04843 | 538 |
| RPS27A;UBA52;UBB;UBC | 1.66 | P62979;P62987;P0CG47;P0CG48 | 48;48;48;48 |
| RPS3A | 1.60 | P61247 | 249 |
| RPS4X | 1.67 | P62701 | 134 |
| RPS7 | 2.62 | P62081 | 142 |
| RSF1 | 2.72 | Q96T23 | 1061 |
| RSF1 | 2.38 | Q96T23 | 1050 |
| RUVBL2 | 2.42 | Q9Y230 | 9 |
| S100A11 | 1.50 | P31949 | 3 |
| SCP2 | 8.24 | P22307 | 453 |
| SDHA | 1.39 | P31040 | 250 |
| SDHA | 1.13 | P31040 | 538 |
| SENP3 | NA | Q9H4L4 | 43 |
| SERBP1 | 1.40 | Q8NC51 | 122 |
| SERBP1 | 1.32 | Q8NC51 | 329 |
| SERBP1 | 0.70 | Q8NC51 | 68 |
| SF1 | 2.57 | Q15637 | 15 |
| SF3B1 | 4.58 | O75533 | 195 |
| SF3B3 | 1.77 | Q15393 | 109 |
| SHMT2 | 1.48 | P34897 | 181 |
| SHMT2 | 1.41 | P34897 | 409 |
| SHMT2 | 1.27 | P34897 | 469 |
| SHMT2 | 1.24 | P34897 | 356 |
| SKP1 | 2.17 | P63208 | 142 |
| SLC25A4 | 1.47 | P12235 | 23 |
| SLC25A5 | 2.94 | P05141 | 199 |
| SLC25A5 | 2.08 | P05141 | 23 |
| SLC25A5;SLC25A31;SLC25A4;SLC25A6 | 2.22 | P05141;Q9H0C2;P12235;P12236 | 92;104;92;92 |
| SLC25A5;SLC25A6 | 2.27 | P05141;P12236 | 96;96 |
| SLC25A6 | 2.22 | P12236 | 105 |
| SLC25A6 | 2.04 | P12236 | 23 |
| SMARCA5 | NA | O60264 | 440 |
| SMC3 | 2.07 | Q9UQE7 | 105 |
| SMC3 | 2.07 | Q9UQE7 | 106 |
| SMCHD1 | 5.12 | A6NHR9 | 771 |
| SMCHD1 | 3.09 | A6NHR9 | 1349 |
| SNRNP200 | 4.54 | O75643 | 46 |
| SOD2 | 1.91 | P04179 | 68 |
| SON | 0.18 | P18583 | 2055 |
| SPTBN1 | NA | Q01082 | 2269 |
| SQRDL | 2.11 | Q9Y6N5 | 135 |
| SUCLG2 | 1.90 | Q96I99 | 338 |
| SUGP2 | 4.39 | Q8IX01 | 1035 |
| SYNRG | 0.41 | Q9UMZ2 | 744 |
| TBL1XR1 | 0.86 | Q9BZK7 | 102 |
| TCOF1 | 2.48 | Q13428 | 155 |
| TCOF1 | 2.27 | Q13428 | 367 |
| TCP1 | 0.95 | P17987 | 400 |
| TECR | 2.77 | Q9NZ01 | 22 |
| TGS1 | 1.18 | Q96RS0 | 510 |
| TGS1 | 1.18 | Q96RS0 | 514 |
| THOC5 | 3.64 | Q13769 | 24 |
| THRAP3 | 3.19 | Q9Y2W1 | 519 |
| TKT | 0.93 | P29401 | 6 |
| TKT | 0.79 | P29401 | 499 |
| TKT | 0.11 | P29401 | 281 |
| TOPAZ1 | 0.95 | Q8N9V7 | 638 |
| TP53BP1 | NA | Q12888 | 1626 |
| TPR | 1.90 | P12270 | 1690 |
| TPR | 1.48 | P12270 | 755 |
| TPR | 1.38 | P12270 | 713 |
| TPR | 1.29 | P12270 | 748 |
| TRIM33 | 38.27 | Q9UPN9 | 951 |
| TRIM33 | 4.27 | Q9UPN9 | 953 |
| TRIM33 | 3.70 | Q9UPN9 | 950 |
| TRMT1L | 1.08 | Q7Z2T5 | 611 |
| TUBA1B;TUBA1C | 1.87 | P68363;Q9BQE3 | 60;60 |
| TUBB4B | 1.24 | P68371 | 58 |
| UBE2V1 | 1.15 | Q13404 | 10 |
| UCHL5 | 1.98 | Q9Y5K5 | 158 |
| UGGT2 | NA | Q9NYU1 | 5 |
| UHRF2 | 2.48 | Q96PU4 | 12 |
| USP49 | 0.83 | Q70CQ1 | 348 |
| VDAC1 | 2.06 | P21796 | 12 |
| VDAC1 | 1.73 | P21796 | 109 |
| VDAC1 | 1.41 | P21796 | 20 |
| VDAC1 | 1.34 | P21796 | 224 |
| VDAC2 | NA | P45880 | 72 |
| VDAC2 | NA | P45880 | 74 |
| VDAC3 | 1.48 | Q9Y277 | 63 |
| WBP11 | 2.41 | Q9Y2W2 | 13 |
| XRCC1 | NA | P18887 | 298 |
| XRCC5 | 1.60 | P13010 | 565 |
| YEATS4 | 2.08 | O95619 | 131 |
| ZMYM2 | 2.77 | Q9UBW7 | 441 |
| ZMYM2 | 2.08 | Q9UBW7 | 1280 |
| ZNF740 | 17.52 | Q8NDX6 | 24 |

**Table S3. Correlation between expression of aldolase A-K13ac and clinico-pathological characteristics in patients with esophageal cancer (n = 258).**

|  |  | **Aldolase A-K13ac** | | |
| --- | --- | --- | --- | --- |
| **Characteristic** | **All cases** | **High** | **Low** | ***P* value** |
| Age |  |  |  | 0.298 |
| < 59 | 134 | 81 (54.7) | 53 (48.2) |  |
| ≥59 | 124 | 67 (45.3) | 57 (51.8) |  |
| Gender |  |  |  | 0.731 |
| Male | 202 | 117 (79.1) | 85 (77.3) |  |
| Female | 56 | 31 (20.9) | 25 (22.7) |  |
| Tumour location |  |  |  | 0.303 |
| Upper thoracic | 12 | 6 (4.1) | 6 (5.5) |  |
| Middle thoracic | 115 | 72 (48.6) | 43 (39.1) |  |
| Lower thoracic | 131 | 70 (47.3) | 61 (55.5) |  |
| Histologic grade |  |  |  | **0.006**** |
| G1 | 31 | 20 (13.5) | 11 (10.0) |  |
| G2 | 204 | 122 (82.4) | 82 (74.5) |  |
| G3 | 23 | 6 (4.1) | 17 (15.5) |  |
| Invasive depth |  |  |  | **0.033*** |
| T1 | 9 | 8 (5.4) | 1 (0.9) |  |
| T2 | 42 | 28 (18.9) | 14 (12.7) |  |
| T3 | 206 | 111 (75.0) | 95 (86.4) |  |
| T4 | 1 | 1 (0.7) | 0 (0.0) |  |
| Lymph node metastasis |  |  |  | 0.415 |
| N0 | 126 | 74 (50.0) | 52 (47.3) |  |
| N1 | 68 | 41 (27.7) | 27 (24.5) |  |
| N2 | 47 | 22 (14.9) | 25 (22.7) |  |
| N3 | 17 | 11 (7.4) | 6 (5.5) |  |
| TNM stage |  |  |  | 0.326 |
| Ⅰ | 14 | 11 (7.4) | 3 (2.7) |  |
| Ⅱ | 116 | 66 (44.6) | 50 (45.5) |  |
| Ⅲ | 111 | 60 (40.5) | 51 (46.4) |  |
| Ⅳ | 17 | 11 (7.4) | 6 (5.5) |  |
| Data are shown as n (%). The *P* value was calculated by calculated by Chi-squared test, unless otherwise stated. | | | | |

**Table S4. Multivariate Cox regression for overall survival and disease-free survival.**

| **Variable** | **Overall survival** | |  | **Disease-free survival** | |
| --- | --- | --- | --- | --- | --- |
|  | **Hazard Ratio (95%CI)** | ***P* value** |  | **Hazard Ratio (95%CI)** | ***P* value** |
| **Aldolase A-K13ac** |  |  |  |  |  |
| High vs. low | 0.57 (0.42 - 0.77) | **< 0.001***** |  | 0.56 (0.42 - 0.76) | **< 0.001***** |
| **Histologic grade** |  |  |  |  |  |
| G3 vs.G2 vs.G1 | 1.45 (1.01 - 2.07) | **0.042*** |  | 1.54 (1.07 - 2.20) | **0.020*** |
| **Invasive depth** |  |  |  |  |  |
| T3+T4 vs. T1+T2 | 1.30 (0.88 - 1.91) | 0.187 |  | 1.35 (0.90 - 2.03) | 0.140 |
| **Lymph node metastasis** |  |  |  |  |  |
| N1+N2+N3 vs. N0 | 1.69 (1.25 - 2.28) | **0.001***** |  | 1.67 (1.24 - 2.26) | **0.001***** |
| Variables were selected with a stepwise selection method. CI denotes confidence interval. | | | | | |

**Table S5. Primers and peptides used in the study.**

| **Primer/Peptide** | **Sequence** |
| --- | --- |
| Mouse WT forward | 5’- GGGGCGTGCTGAGCCAGACCTCCAT-3’ |
| Mouse WT reverse | 5’-TCCCGACAAAACCGAAAATCTGTGG-3’ |
| Mouse L2Δ13 forward | 5’- TGCATCGCATTGTCTGAGTAGG-3’ |
| Mouse L2Δ13 reverse | 5’-TCCCGACAAAACCGAAAATCTGTGG-3’ |
| Human ALDOA-K13R forward | 5’- ACTGACCCCGGAGCAGAGGAAGGAGCTGT-3’ |
| Human ALDOA-K13R reverse | 5’-CTCTGCTCCGGGGTCAGTGCTGGATATTGG-3’ |
| Human ALDOA-K13Q forward | 5’- ACTGACCCCGGAGCAGCAGAAGGAGCTGT-3’ |
| Human ALDOA-K13Q reverse | 5’-GCTGCTCCGGGGTCAGTGCTGGATATTGG-3’ |
| Aldolase A-K13ac peptide 1 | ALTPEQK (ac) KELSDIC |
| Aldolase A-K13ac peptide 2 | YPALTPEQK (ac) KELSDC |

**Table S6. Antibodies used in the study.**

| **Antibody** | **Company** | **Catalog Number** | **Dilution** | **Applications** |
| --- | --- | --- | --- | --- |
| Acetylated-Lysine (AcK) | PTM Biolabs Inc | PTM-102 | 1:100 | IP |
| Acetylated-Lysine (AcK) | CST | 9441S | 1:1000 | WB |
| Actin-stain 555 phalloidin | Cytoskeleton | PHDH1 | 1:200 | IF |
| Akt | CST | 9272S | 1:1000 | WB |
| Aldolase A | Abcam | ab54770 | 1:1000 | WB |
| Aldolase A | Santa Cruz Biotechnology | sc-390733 | 1:200 | IF |
| Aldolase A-K13ac | PTM Biolabs Inc | CTM-205 | 1:10000; 1:4000 | WB; IHC |
| α Enolase | Santa Cruz Biotechnology | sc-101513 | 1:1000 | WB |
| α-SMA | Abcam | ab5694 | 1:500 | IHC |
| β-actin | Santa Cruz Biotechnology | sc-47778 | 1:1000 | WB |
| DAPI | Sigma-Aldrich | D9564 | 0.1 μg/mL | IF |
| Flag (DDDDK tag) | Proteintech | 20543-1-AP | 1:100 | IHC |
| Flag (M2) | Sigma-Aldrich | F3165 | 1:5000 | WB; IP |
| FLAG M2 Affinity Gel | Sigma-Aldrich | A2220 | - | IP |
| FLAG Peptide | Sigma-Aldrich | F3290 | - | IP |
| GAPDH | Santa Cruz Biotechnology | sc-47724 | 1:1000 | WB |
| GAPDH | Sigma-Aldrich | G9545 | 1:100 | IF |
| HA | Santa Cruz Biotechnology | sc-7392 | 1:1000 | WB; IP |
| GST | TransGen Biotech | HT601 | 1:5000 | WB |
| GST Bind Resin | Millipore | 70541 | - | IP |
| ki67 | ZsBio | ZM-0166 | prediluted | IHC |
| L2Δ13 | Bioss | - | 1:100; 1:500 | IF; IHC |
| LOXL2 | Abcam | ab96233 | 1:1000 | WB |
| LOXL2 | Novus | NBP1-32954 | 1:100; 1:500 | IF; IHC |
| normal rabbit IgG | Santa Cruz Biotechnology | sc-2763 | 1:1000 | WB |
| normal mouse IgG | Santa Cruz Biotechnology | sc-2762 | 1:1000 | WB |
| p-Akt (S473) | CST | 9271S | 1:1000 | WB |
| TPI | Santa Cruz Biotechnology | sc-100541 | 1:1000 | WB |
| Vimentin | Millipore | CS207806 | 1:500 | WB |
